# *Schistosoma mansoni* cercarial invadolysin cleaves complement C3 and suppresses proinflammatory cytokine production

**DOI:** 10.1101/2022.09.16.508213

**Authors:** Jacob R. Hambrook, Patrick C. Hanington

## Abstract

*Schistosoma mansoni* employs immune evasion and immunosuppression to overcome immune responses mounted by its snail and human hosts. Myriad immunomodulating factors underlie this process, some of which are proteases. Here, we demonstrate that one protease, an invadolysin we have termed SmCI-1, is released from the acetabular glands of *S. mansoni* cercaria and is involved in creating an immunological milieu favorable for survival of the parasite. We confirm the presence of SmCI-1 in the cercarial stage of *S. mansoni* and demonstrate that it is released during transformation into the schistosomula. We confirm SmCI-1 functions as a metalloprotease with the capacity to cleave collagen type IV, gelatin and fibrinogen. We also show that complement component C3b is cleaved by this protease, resulting in inhibition of the classical and alternative complement pathways. Finally, we assess the effect of SmCI-1 on cytokine release from human peripheral blood mononuclear cells, providing compelling evidence that SmCI-1 promotes an anti-inflammatory microenvironment by enhancing production of IL-10 and suppressing the production of inflammatory cytokines like IL-1B and IL-12p70 and those involved in eosinophil recruitment and activation, like eotaxin-1 and IL-5.

## Introduction

Despite significant investment into treatment and prevention programs, schistosomiasis continues to afflict an estimated 230 million people worldwide (1). Key among the numerous ways in which schistosomes persist in human populations, in spite of efforts to eliminate the parasites, are immunomodulatory mechanisms that interfere with and suppress the immune response of their gastropod and human hosts (2–4). While the strategies used to overcome immune responses are highly effective, they are not perfect. Humans can develop low levels of non-sterilizing immunity to infection, with exposure to adult worm antigens being a likely mechanism (5,6). Lung stage schistosomula are also a possible target for immune mediated destruction given their ability to elicit a strong immune response in murine models, as well as the discovery that plasma from self-curing Rhesus macaques is capable of enhanced killing of 3-5 day old schistosomula (7,8). Such findings demonstrate that an in-depth examination of the survival mechanisms at various stages of the intra-human schistosome life cycle merits investigation as possible targets for novel therapeutics and/or vaccine development efforts.

Proteases are prominently featured amongst the immunomodulatory mechanisms employed by *Schistosoma mansoni* during the snail and human infection process (9–11). In snails, *S. mansoni* miracidia have been shown to release calpain, serine peptidases, and at least one metallopeptidase (12,13). Gene array data from developing sporocysts also feature proteases such as cathepsins, elastase, and hemoglobinase, each of which are upregulated during development within the snail (10,11). In humans, the employment of proteases by cercaria is widespread and well-studied (14,15). In contrast, the use of proteases and immune modulators remains less extensively examined in lung-stage schistosomula due to the relative difficulty of culturing and isolation this stage of the parasite (3). Adult schistosome-produced proteases that have been examined to date are largely involved in anti-coagulation mechanisms (16–18). Other proteases, such as metalloproteases and aminopeptidases, have been identified in adult worm extracts, but remain to be functionally characterized (9).

Proteases are especially important during the two tissue-penetrative stages of the life cycle of *S. mansoni*. These enzymes have been most extensively studied within the context of cercarial penetration of, and residence within human skin. During this time, the developing parasites strive to locate a venule through which they enter the circulation (15,19,20). Upon attachment to the surface of the skin, cercaria begin migrating through the numerous layers of the epidermis, a process involving mechanical disruption of host cell-to-cell interactions as well as the release of proteases from the acetabular glands (15,21). In *S. mansoni*, proteases released by the acetabular glands are predominantly serine proteases, which have been demonstrated to cleave key structural components in skin such as elastin, laminin, fibronectin, collagen type IV, and keratin (22–25). So crucial are these serine proteases that one, a 28/30kda protease termed *S. mansoni* cercaria elastase (SmCE) makes up roughly 36% of the contents of the acetabular glands (26). Broad inhibition of serine proteases, as well as SmCE specifically, substantially reduces cercarial penetration success, highlighting the essential function of SmCE during the initial stages of *S. mansoni* infection of their human host (25,27). Cleavage targets of this serine protease are not limited to structural host components, with SmCE having been shown to cleave complement component C3 (22). This highlights the reality that cercaria must not only combat the structural components of human skin, but also encounter immunological factors. The capacity of SmCE to cleave complement component C3 suggests that cercaria have developed ways to circumnavigate the immune factors present in the skin that may target them for destruction. Accompanying these serine proteases are cysteine proteases, which are less well understood in the context of *S. mansoni*, but which are 40X more abundant in *S. japonicum* cercaria (28,29). Also present are several metalloproteases that have yet to be functionally characterized (26,30).

The *S. mansoni* genome features 114 annotated metalloproteases, 35 of which have been found to be differentially transcribed during distinct parasite life cycle stages (31,32). Key among the metalloproteases found in *S. mansoni* cercaria are a subset of matrix metalloproteases (MMPs) known as invadolysins. Invadolysins, like all MMPs, employ a zinc ion, coordinated by histidine residues, in order to attract and retain water molecules to facilitate proteolysis (33,34). They are typically synthesized as zymogens, awaiting the removal of an inhibitory-cysteine-containing pre-protein fragment in order to be rendered active (35). As their name suggests, the proteolysis performed by MMPs frequently target the extracellular matrix (ECM) components of cells such as collagen, laminin, and elastin (33,36). Invadolysins are considered M8 metzincin metalloproteases, which employ a methionine in a characteristic “met-turn” involved in the stability of the protease, and from which the name metzincin is derived (37,38). Seven invadolysins have been shown to be differentially regulated in *S. mansoni* to date, with five being most highly transcribed in the germball stage of development, during which the necessary products for skin penetration by cercaria are generated (32). The other two are most highly upregulated in cercaria themselves (32). Invadolysins are also found in *S. japonicum* cercaria, with the three upregulated invadolysin genes observed in an *S. japonicum* micro-array bearing high levels of amino acid similarity to three of the most highly upregulated invadolysins in *S. mansoni* (39). The identification of these invadolysins via transcriptomics analysis, as well as their presence in both *S. mansoni* and *S. japonicum*, render them interesting targets for further study. The case for characterization of invadolysins during initial infection of a human host is bolstered by the observation that the second most highly upregulated invadolysin in *S. mansoni* germballs (Smp_090100.1) has also been identified as the second most abundant protein released from *S. mansoni* acetabular glands during transformation into schistosomula, comprising roughly 12.8% of the normalized volume of the acetabular glands (26). For the purposes of this work, we refer to Smp_090100.1 as *S. mansoni* cercarial invadolysin 1 (SmCI-1) in keeping with the nomenclature surrounding cercarial elastase (SmCE).

The utilization of invadolysins as tools for invasion of a host by a parasite has been extensively studied, especially in the case of kinetoplastid parasites such as *Leishmania spp*. and *Trypanosoma spp*. Invadolysins were initially characterized in *Leishmania spp*. and are often termed leishmanolysins, or glycoprotein 63kda (GP63) (40,41). GP63 can effectively cleave gelatin, albumin, fibrinogen and collagen type IV (42,43). McGwire et al. demonstrate that the capacity for GP63 to degrade key ECM proteins is involved in facilitating leishmania movement through a Matrigel coated membrane, implying that GP63 can aid leishmania in movement throughout its host and could help promastigotes penetrate mononuclear phagocytic cells (42). In addition to structural molecules, *Leishmania major* GP63 can cleave IgG and CD4, and facilitate the conversion of C3b into iC3b, thereby increasing uptake of the parasite into host immune cells (44–49). GP63 also prevents complement mediated lysis of promastigotes in human serum (45). Once inside of a host macrophage, GP63 prevents degradation of proteins by the phagolysosome, while also altering intracellular signalling pathways by cleaving the myristoylated alanine-rich kinase substrate (MARKS)-related protein (MRP), as well as NF-κB (50–52). In addition to these functions, GP63 is capable of reducing chemotactic responses in cells such as macrophages, neutrophils and monocytes (53,54). The ability for neutrophils and monocytes to initiate a respiratory burst response in response to opsonized zymosan is also hampered by the presence of *L. major* GP63 (54) To add to all of these functions, GP63 is also capable of reducing NK cell proliferation (55).

To our knowledge, the only helminth metalloprotease to have been functionally characterized as a known invadolysin was our previous work on a protein termed SmLeish (Smp_135530) that was shown to inhibit the movement of *Biomphalaria glabrata* haemocytes, and to negatively impact the kinetics of *S. mansoni* establishment within *B. glabrata* (13). Other helminth metalloproteases have been implicated in immunomodulation as well, with *Necator americanus* metalloproteases demonstrating a specific anti-eosinophil mechanism of action via the cleaving of eotaxin-1, but not IL-8 (56).

Given the consistent immunomodulatory role played by invadolysins during parasite infection/establishment, and the presence of SmCI-1 as the second most abundant protein in *S. mansoni* acetabular glands, we set out to characterize SmCI-1 in the context of early *S. mansoni* infection of the human host. In this study, we demonstrate that SmCI-1 acts as a MMP and can cleave specific human proteins relevant to structural and immunological functions. Additionally, we examined the capacity of SmCI-1 to alter cytokine output of human polymorphic blood mononuclear cells. This assessment identified anti-inflammatory properties. To follow up on this observation we used a mutant form of SmCI-1 lacking MMP activity to demonstrate that reduction of key inflammatory cytokines required MMP activity, while eliciting IL-10 production did not. All of this was done with the intent of better understanding how *S. mansoni* is capable of surviving within human skin, as well as gaining a deeper knowledge of how parasitic organisms employ metalloproteases during infection and establishment within a host.

## Methods

### Structure predictions and visualization

Homology modeling was performed by submitting the amino acid sequence for SmCI-1 (Smp_090100.1) to the Robetta Server (robetta.bakerlab.org). RoseTTAFold, a deep learning-based modeling method, was used for secondary/tertiary structure prediction. The prediction with the highest confidence (0.71) was uploaded to PyMol (PyMOL™ Version 2.4.1 Schrodinger ©) for visualization and labeling of key active site amino acids. For GP63 comparison, the GP63 protein databank file was obtained from the RCSB protein databank (57).

### Production of Recombinant SmCI-1 and Purification

The genetic sequence of SmCI-1 was synthesized and inserted into a pHEK293 Ultra Expression Vector I. Transient transfection of HEK293 cells was performed, with the recombinant protein being purified via the anti-SmCI-1 polyclonal antibody generated against a recombinant SmCI-1 protein generated in a bacterial expression system (see below). A mutant form of SmCI-1 (MutSmCI-1) was synthesized in an identical manner, with the Glu232 codon (GAA) replaced with a Glycine codon (GGG).

### Production of Rabbit anti-SmCI-1 Antibody

The synthesized Acetabular genetic sequence was inserted into the pET-47b(+) bacterial expression vector (Novagen) following manufacturers specifications. BL21(DE3) (ThermoFisher Scientific) cells were transfected with SmCI-1-containing pET-47b(+) vector, grown at 37°C in LB media (ThermoFisher Scientific) containing 100μg/ml Kanamycin (ThermoFisher Scientific) and then induced with 1 mM isopropyl β-d-1-thiogalactopyranoside (IPTG) once the cells had grown to OD 600. Following ∼4 hours of growth, the cells were concentrated by centrifugation at 10 000 rpm for 20 minutes at 4°C. The final weight of the bacteria pellet was measured and then the lysing reagent B-PER (Thermo Fisher Scientific) was added at a concentration of 4 ml/g of bacteria in combination with phenylmethylsufonyl fluoride (PMSF, final concentration of 1 mM), mixed gently and left to incubate at room temperature for 15 minutes. Before application to the 5ml 6xHIS column (GE Healthcare) for purification, the supernatant was diluted to a total protein concentration of 100 μg/ml in binding buffer (GE Healthcare). Following purification, the 6x HIS tag region of the recombinant protein was removed following the vector manufacturers specifications (Novagen). This recombinant SmCI-1 was used to generate a rabbit anti-Sm-CI-1 polyclonal antibody using the Custom pAb service offered by Genscript.

### *S. mansoni* Culture

M-line *Biomphalaria glabrata* snails were experimentally infected with 10 *Schistosoma mansoni* (NMRI strain) miracidia in 12 well plates for 24 hours, before being raised in artificial spring water for 6-8 weeks, at room temperature, on a diet consisting of red leaf lettuce. Snails were allocated into 6 well plates and exposed to light for 5 hours during which cercaria emerged. Cercaria were mechanically transformed into schistosomula by vortexing in 1.5 ml tubes.

Parasites were kept on ice for 15 minutes, before being centrifuged for 5 minutes at 1000g. Artificial spring water was removed, and 200 μl of RPMI 1640 was added. Cercaria were again vortexed for one minute, and visually inspected using microscopy to confirm detachment of their tails. Parasites were centrifuged again, supernatants were collected and frozen for western blotting, and 200 μl of RPMI 1640 (1% Penicillin/Streptomycin) was again added to the transformed schistosomula. Schistosomula were cultured at 37°C for 24 hours. Untransformed cercaria, transformed schistosomula, and schistosomula excreted/secreted (E/S) products were all gathered for western blot analysis.

### Western blotting

Parasite samples were resuspended with PBS, to which 4X reducing Laemmli buffer was added, before being boiled at 95°C for 15 minutes. Samples were loaded into 12% SDS PAGE gels with a final concentration of ∼20 parasites worth of material per lane. Samples were run on the Mini PROTEAN Tetra system (Bio-Rad) at 150 V and 180 mA, and then blotted for 2 hours onto 0.45 μm supported nitrocellulose membranes (Bio-Rad). Membranes were blocked for one hour using 5% Bovine Serum Albumin (BSA) in tris buffered saline with 0.1% Tween-20 (TBS-T) at room temperature on a rocking platform. A 1:1000 dilution of the rabbit anti-SmCI-1 antibody was added for an hour. The membranes were washed 1X 10 min with TBS-T, 2X 5 min with TBS-T, and 1X 5 min with TBS. An HRP-conjugated goat anti-rabbit polyclonal secondary antibody was then diluted 1:1000 in 5% BSA in TBS-T and applied to the membrane for 1 hour. Membranes were then washed as previously described. Development of the membranes was done using SuperSignal™ West Dura Extended Duration Substrate (Thermo Scientific), and chemiluminescent results were visualized using an ImageQuant LAS 4000 Gel Imaging System (GE Healthcare).

### Immunofluorescence staining

Freshly shed cercariae were fixed using 4% paraformaldehyde for 1 hour at room temperature in 1.5 ml tubes. Fixed cercariae were washed 2X using 1000 μl Phosphate buffered saline + 0.1% Tween-20 (PBS-T). Permeabilization was performed using 500 μl of PBS + 5%BSA, 0.02% NaN_3_, 1% TritonX-100 for 1 hour at room temperature. Cercariae were then washed 2X as described above. Parasites were blocked for 1 hour at room temperature in PBS +5% BSA after which a 1:400 dilution of rabbit anti-SmCI-1 was added for 1 hour. Samples were washed 2X 20 minutes in PBS-T on a rocking platform, and then stained with a 1:400 dilution of donkey anti-rabbit Alexa 568 antibody and 10 μl of Alexa Fluor™ 488 Phalloidin (ThermoFisher Scientific) at 4°C overnight. Samples were again washed 2X 20 minutes with 1000 μl PBS-T and 2X 20 minutes with 1000 μl PBS. Finally, samples were resuspended in 100 μl of PBS, to which one drop of Fluoroshield™ with DAPI (Millipore Sigma) was added. Fluorescence was visualized using a Leica TCS SP5 laser scanning confocal microscope before analysis via ImageJ (NIH).

### Generic MMP assay

Generic MMP function was determined using a Generic MMP Activity Kit (AnaSpec, AS-72202) as per the kit’s instructions. Recombinant SmCI-1 and MutSmCI-1 at concentrations of 0.25, 0.5, and 1.0 μg/ml were added to a kit specific reaction buffer (KSRB) and were either activated via incubation with 1mM 4-Aminophenylmercuric acetate (APMA) for one hour at 37°C or left at 37°C without activation. Inhibition was assessed via the addition of 250 μM of 1,10-phenanthroline (Sigma Aldrich) prior to addition of the substrate. As a positive control, recombinant human MMP-8 (R&D Systems) was subjected to the same conditions. Substrate only, buffer only, and APMA controls were also added. 45 μl samples were then mixed with 45 μl of substrate (Mca/Dnp fluorescence resonance energy transfer peptide) in a clear-bottom black-welled 96 well plate and 330nm/390nm fluorescence readings were obtained after 1 hour using a SpectraMax M2 microplate reader (Molecular Devices). Difference between treatments were assessed via One-Way Anova.

An additional experiment was run to determine the activity levels of SmCI-1 in various buffers of relevance to other assays. Generic MMP activity in KSRB, phosphate buffered saline (PBS, PH 7.4), RPMI 1640 (Gibco), DMEM (Gibco), Krebs-Ringer Phosphate Buffer (KRPG, PH7.4), and MMP buffer (50mM Tris, 10mM CaCl_2_, 150mM NaCl, PH 7.5) was examined after 3 hours of incubation at 37°C.

### Gelatin, collagen and elastin cleavage assay

Where possible, the capacity of SmCI-1 to cleave structural molecules was assessed using commercially available kits. Due to the variability of activity between buffers, we elected to use the KSRB from the Generic MMP Activity Kit for these assays. All MMPs were activated via incubation with 1mM APMA as previously described. Gelatin and collagen type 4 cleavage were examined using an EnzChek™ Gelatinase/Collagenase Assay Kit (ThermoFisher Scientific), with MMP-2 (AnaSpec) and *Clostridium histolyticum* collagenase (ThermoFisher Scientific) used as positive controls. Proteases were incubated with 100 μg/ml fluorescein labeled substrate in a clear-bottom black-welled 96 well plate and 495nm/515nm fluorescence readings were obtained after 20 hours incubations at 37°C using a SpectraMax M2 microplate reader (Molecular Devices). Difference between treatments were assessed via One-Way Anova.

Elastin cleavage was examined using a SensoLyte® Green Elastase Assay Kit (AnaSpec). Porcine elastase (AnaSpec) and MMP-12 (R&D Systems) were used as positive controls. Cleavage of the 5-FAM/QXL™ 520 labeled elastin was measured in a clear-bottom black-welled 96 well plate and 490nm/520nm fluorescence readings were obtained after 1 hour at 37°C using a SpectraMax M2 microplate reader (Molecular Devices). Difference between treatments were assessed via One-Way Anova.

### Silver stain cleavage assays

For relevant structural and immunological proteins for whom no relevant cleavage assay was found, silver staining was used to determine the capacity of SmCI-1 to cleave such molecules. Two micrograms of Human Fibrinogen (Sigma-Aldrich), Immunoglobulin G (Sigma-Aldrich), recombinant complement component C3 (Millipore Sigma), recombinant CD4 (Sigma Aldrich) were added to 20 μl of KSRB with 1mM APMA. Samples were then either left without protease or given 2 μg/ml SmCI-1, or 2 μg/ml Trypsin as a positive control for cleavage.

Samples were incubated at 37°C for 18 hours, before being treated with reducing Laemmli buffer, boiled at 95°C for 15 minutes, and run on 12% and 15% SDS PAGE gels using a Mini PROTEAN Tetra system (Bio-Rad) at 150 V and 180 mA. Silver staining was performed using a Pierce™ Silver Stain Kit (ThermoFisher Scientific) and images were obtained using an ImageQuant LAS 4000 Gel Imaging System (GE Healthcare).

#### Testing hemolytic activity of human sera pre-treated with rSmCI-1 and rSm-CI-1(Mut)

Human sera (Sigma Millipore; S7023) was preincubated with rSmCI-1 or rSmCI-1Mut at a concentration of 2μg/ml for 18 hours. Using pre-sensitized sheep erythrocytes(ssRBC) (CompTech, USA; B202), classical pathway mediated hemolysis was assessed following previously published protocols (58). Briefly, ssRBCs were washed 1x in veronal buffered saline (VBS) (5 mM Veronal, 145 mM NaCl, 0.15 mM CaCl_2_ 0.5 mM MgCl_2_. 0.025% NaN_3,_ pH 7.3) by centrifugation at 2000 g for 10 minutes at 4°C and then resuspended to 10^9^ cells/ml in 5ml of gelatin veronal buffer (GVB) (VBS + 0.1% gelatin). The ssRBCs were then warmed to 37°C in a water bath. The experimental human sera control and sera with rSmCI-1 or rSmCI-1Mut added was diluted 1:50 in and then further diluted 1:3 prior to addition of 50 μl to a round bottom 96-well plate. Then 50 μl of the ssRBC suspension was added to the wells. Each treatment was replicated 8x on the plate, including a 100% lysis control which contained 50 μl ssRBCs and a cell blank which contained 50μl ssRBCs with 50 μl GVB. Plates were incubated at 37°C for 30 minutes with gentle agitation. Then 150 μl of ice-cold GVB was added to all experimental and cell-blank control wells and 200 μl of distilled water was added to the 100% lysis control wells. The plate was centrifuged at 1000 g for 5 minutes prior to the transfer of 200 μl of supernatant from each well to a new flat-bottomed 96-well plate. Plates were read using spectrophotometer at 412 nm, the cell blank absorbance was subtracted from each measurement to obtain correct absorbances. Fractional hemolysis in each well was calculated relative to the 100% lysis wells.

Alternative pathway-mediated hemolysis was assessed using rabbit erythrocytes (Innovative Research, USA; IRBRBC10ML) following a previously published protocol (58). Briefly, a 1:25 dilution of the SmCI-1 and SmCI-1Mut preincubated sera prepared as described above was prepared and then diluted 1:3 to create a working solution that was added to the wells of a round-bottom 96-well plate. 50 μl of the working sera solution was added along with 50 μl of rabbit erythrocytes at a concentration of 10^8^ cells per ml suspended in AP buffer (GVB containing 5 m*M* Mg2+ and 5 m*M* ethylene glycol-bis[β-aminoethyl ether]N,N’-tetraacetic acid (EGTA)). Each treatment was replicated 8x on the plate, including a 100% lysis control which contained 50 μl rRBCs in AP buffer and a cell blank which contained 50 μl rRBCs with 50 μl AP buffer. Plates were incubated at 37°C for 30 minutes with gentle agitation. Then 150 μl of ice-cold N-saline (9 g NaCl dissolved in 1 L water) was added to all experimental and cell-blank control wells and 200 μl of distilled water was added to the 100% lysis control wells. The plate was centrifuged at 1000 g for 5 minutes prior to the transfer of 200 μl of supernatant from each well to a new flat-bottomed 96-well plate. Plates were read using spectrophotometer at 412 nm. Fractional hemolysis in each well was calculated relative to the 100% lysis wells.

### Cytokine levels

Investigations into the effect of SmCI-1 and MutSmCI-1 on cytokine production were examined using a Human XL Cytokine Luminex Performance Panel Premixed Kit (bio-techne/R&D systems). Human blood mononuclear cells (PBMCs) (Cedarlane/ATCC) were cultured in RPMI at 10,000 cells per treatment. Cells were either left unstimulated, stimulated with 5 cercaria worth of whole cercarial lysate (WCL), or stimulated with 1μg/ml of lipopolysaccharide (LPS). WCL had been generated via 3x 15s sonification of freshly emerged cercaria in PBS, on ice. The effects of the recombinants on cytokine production were assessed by the addition of rSmCI-1 or rMutSmCI-1 at a concentration of 2μg/ml 30 minutes prior to stimulation. Cytokine levels were measured 24 hours post stimulation.

#### Western blot cytokine cleavage assay

Recombinant human Eotaxin-1 (R&D systems) and recombinant IL-5 (R&D systems) at an amount of 50 ng/sample were incubated overnight in kit specific reaction buffer with 1μg/ml rSmCI-1, 1 μg/ml rMutSmCI-1 or 1 μg/ml trypsin. Western blotting was performed as detailed above, using a 1:1000 dilution of monoclonal mouse anti-Eotaxin-1 or IL-5(R&D).

### Statistical analysis

Differences in activity in all cleavage assays, as well as the difference between treatments in our cytokine activity assays were each assessed via One-Way Anova.

## Results

### Predicted Structure of SmCI-1

Modelling the 3D structure of SmCI-1 demonstrated consistency with the core features found in invadolysins (Figure 1). SmCI-1 was predicted to feature an active site, similar to GP63, with three histidines (His231, His235, and His336) present in order to coordinate the immobilization of a zinc molecule, and a glutamic acid (Glu232) utilized as a nucleophile during proteolysis. SmCI-1 also featured amino acids in locations likely to correspond to canonical features of invadolysins, such as the methionine (Met347) needed for a met-turn, and three cysteines (Cys182, Cys194, and Cys211) on the N-terminal side of the active site, each of which is a candidate for a cysteine-switch activation mechanism.

**Figure 1.**
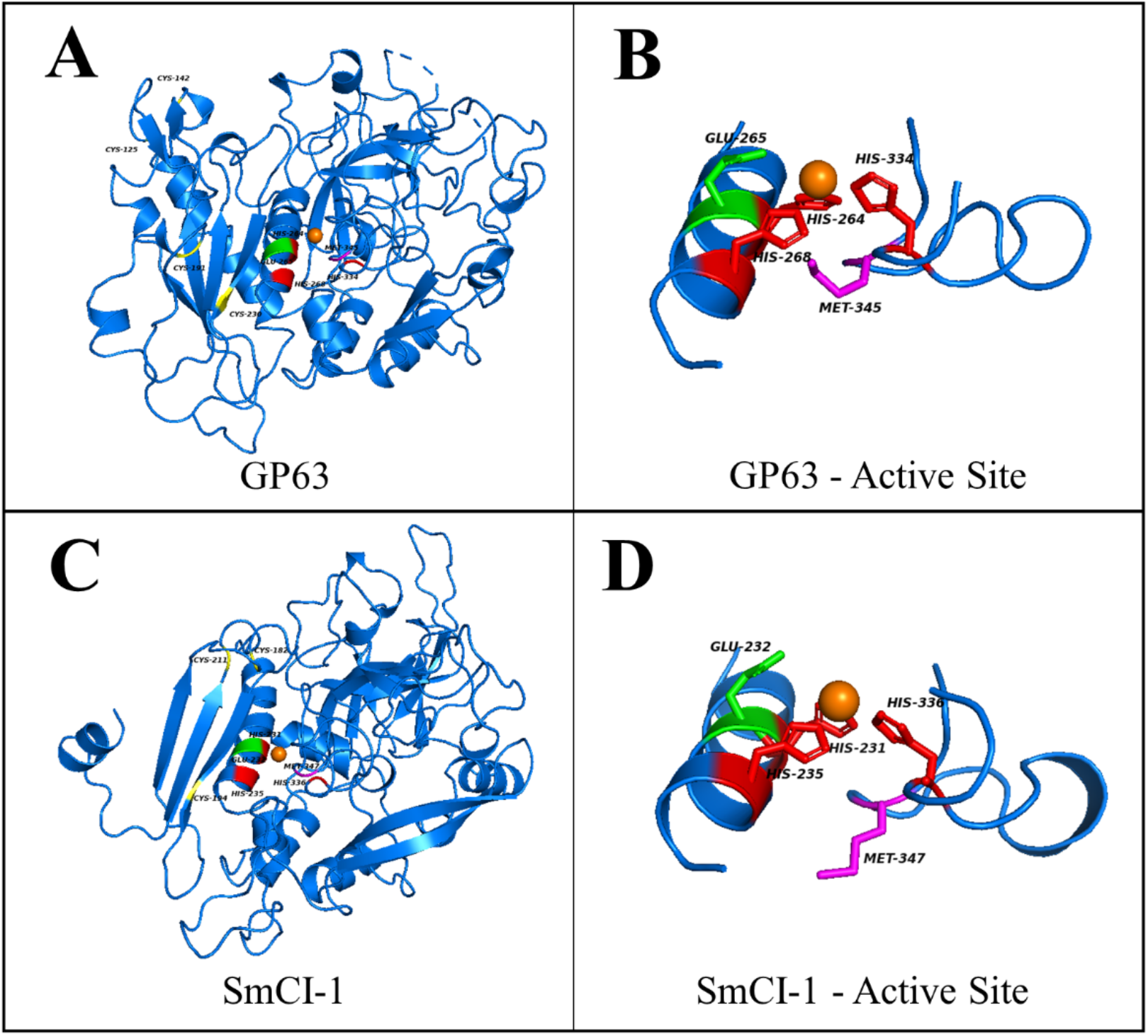
A 3D visualization of the crystal structure of GP63 (A), and a magnified view of its active site (B). The predicted structure of the full amino acid sequence of SmCI-1 is also present (C) alongside a magnified view of its active site (D). Active site histidines are presented in red, holding a zinc ion (orange ball) in position. Nucleophile Glutamic acids are marked as green, with likely met-turn methionines marked as purple. All Cysteines N-terminal to the active site have been coloured yellow.

### Location and expulsion of SmCI-1

Immunofluorescence staining utilizing our polyclonal anti-SmCI-1 antibody revealed a concentrated fluorescence signal emanating from the acetabular glands of *S. mansoni* cercaria (Figure 2A). Staining with either primary or secondary antibodies only failed to elicit such a signal. Western blot analysis of SmCI-1 release during transformation into schistosomula further confirmed the acetabular gland localization (Figure 2B). Our antibody recognizes a protein appearing at the expected size of 65kDa. The 65kDa band was seen in untransformed cercaria, as well as 24 hr post transformation cercaria and their E/S products.

**Figure 2.**
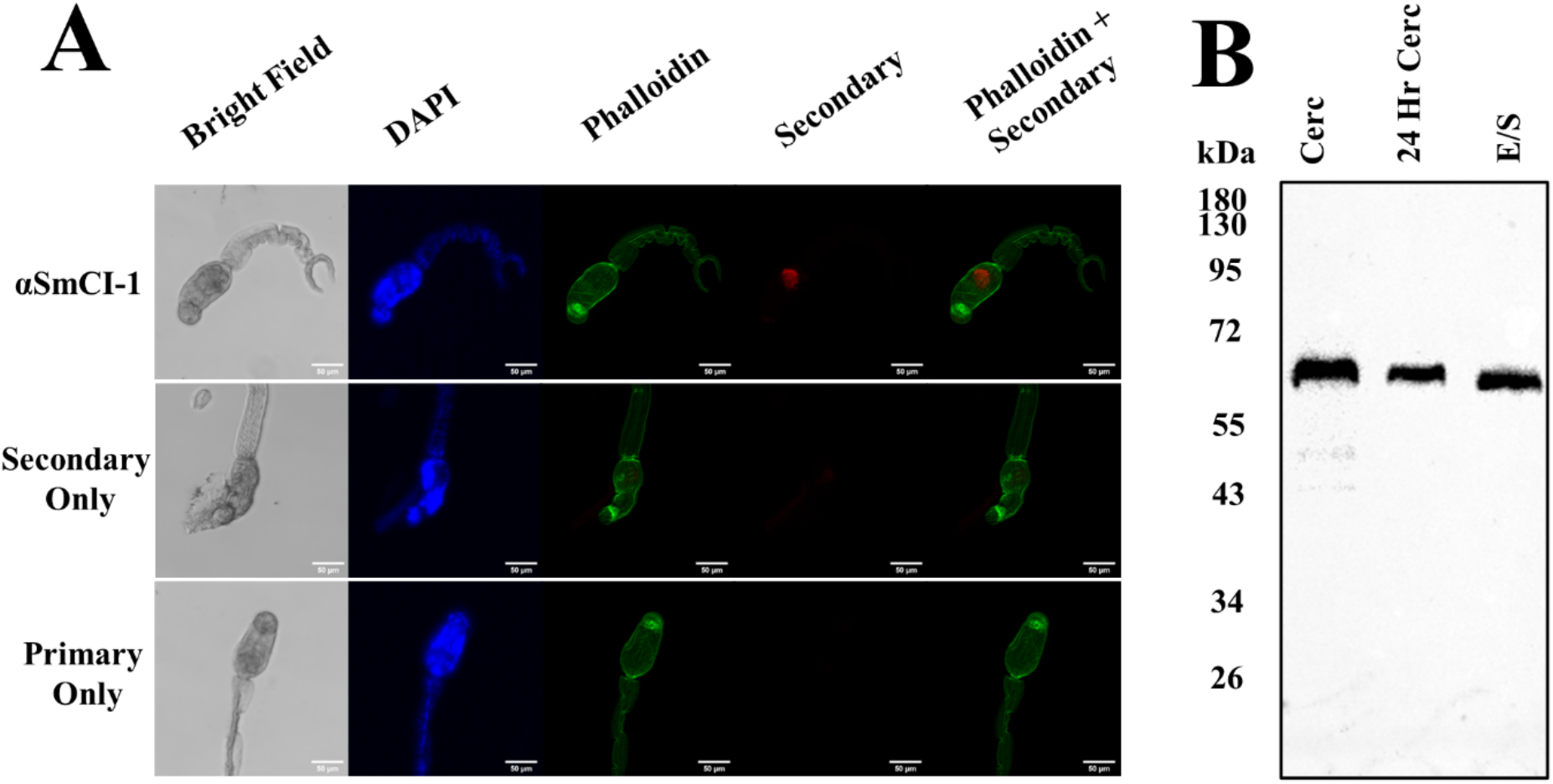
(A) SmCI-1 is localized to the acetabular glands of *S. mansoni* cercaria, as revealed by immunofluorescence imaging, with SmCI-1 signal appearing in red. (B) SmCI-1 is expelled from the cercaria during the initial stages of maturation into schistosomula that occur in human skin. SmCI-1 is shown via western blotting as a ∼65kDa band in cercaria as well as cercarial E/S products.

#### SmCI-1 displays MMP activity

Recombinant SmCI-1 displays dose-dependent MMP activity and is inhibited partially by the addition of [50μM] 1,10-phenanathroline, with activity being completely abrogated via the addition of [250μM] of 1,10-phenanthroline (Figure 2A). Samples left inactivated, and those of the same concentration that were activated with 1mM APMA did not differ significantly in their activity. The mutation of Glu232→Gly232 rendered the rSmCI-1 mutant completely inactive, with no concentration up to and including 1μg/ml differing from the vehicle control with respect to general MMP function (Figure 2B). The positive control, human MMP8 also demonstrated MMP activity capable of being inhibited by the addition of 1,10-phenanthroline (Figure 2C). Unlike rSmCI-1, MMP8 samples activated with APMA had significantly higher MMP activity than those that were not activated. Among MMP8 samples that were not activated, only those at 1μg/ml displayed activity levels that differed significantly from the vehicle control.

#### Cleavage of Structural Substrates

To better understand the role of SmCI-1 in degradation of prominent components of the extracellular matrix of host skin cells and the basement membrane of human skin, we examine the capacity of SmCI-1 to cleave gelatin, collagen type IV, and elastin. In the case of each of these possible substrates, rSmCI-1 was able to weakly cleave each of them. In the cases of gelatin (Figure 4A) and collagen type IV (Figure 4B), rSmCI-1 cleavage activity significantly differed from the APMA negative control, but only at 2μg/ml, and not 1μg/ml. This activity was significantly lower than that seen in the positive control, *Clostridium histolytica* collagenase. In the case of elastin, rSmCI-1 displayed cleavage activity above that of the APMA control for both 2μg/ml and 1μg/ml treatments. This activity was again lower than established positive controls for the cleavage of elastin, such as porcine elastase and human MMP12 (Figure 4C).

**Figure 3.**
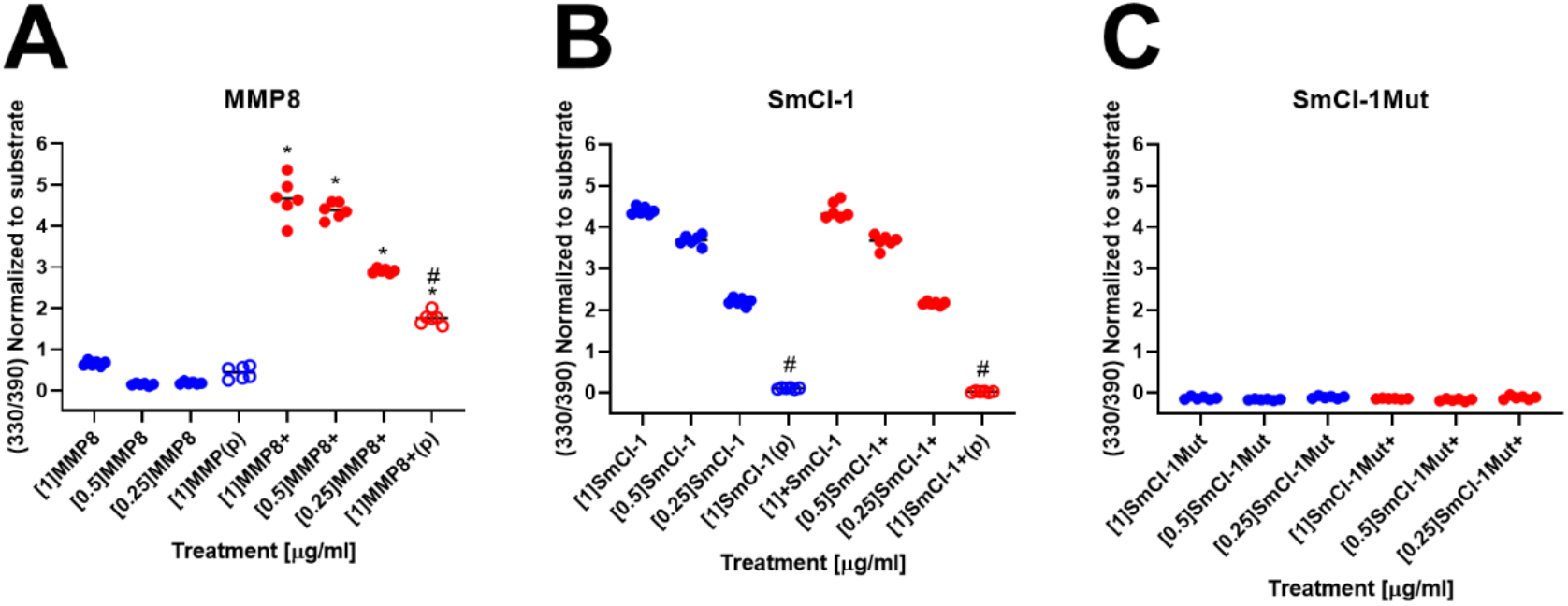
Recombinant SmCI-1 is a functional matrix metalloprotease. MMP activity of (A) human MMP8, (B) rSmCI-1, and (C) rMutSmCI-1. All proteases were examined at three initial concentrations of 1μg/ml, 0.5μg/ml, and 0.25μg/ml. Whereas MMP8 and rSmCI-1 demonstrated activity, rMutSmCI-1 did not. Human MMP8 required activation with 1mM APMA(+), rSmCI-1 had similar activity levels with and without APMA-mediated activation. Significant inhibition via the addition of [50μM] (p) or [250μM] (P) of 1,10-phenanthroine is denoted by (#). Activated samples (+) that differ significantly from their non-activated counterparts of an identical concentration are denoted using (*). (n=6)

**Figure 4.**
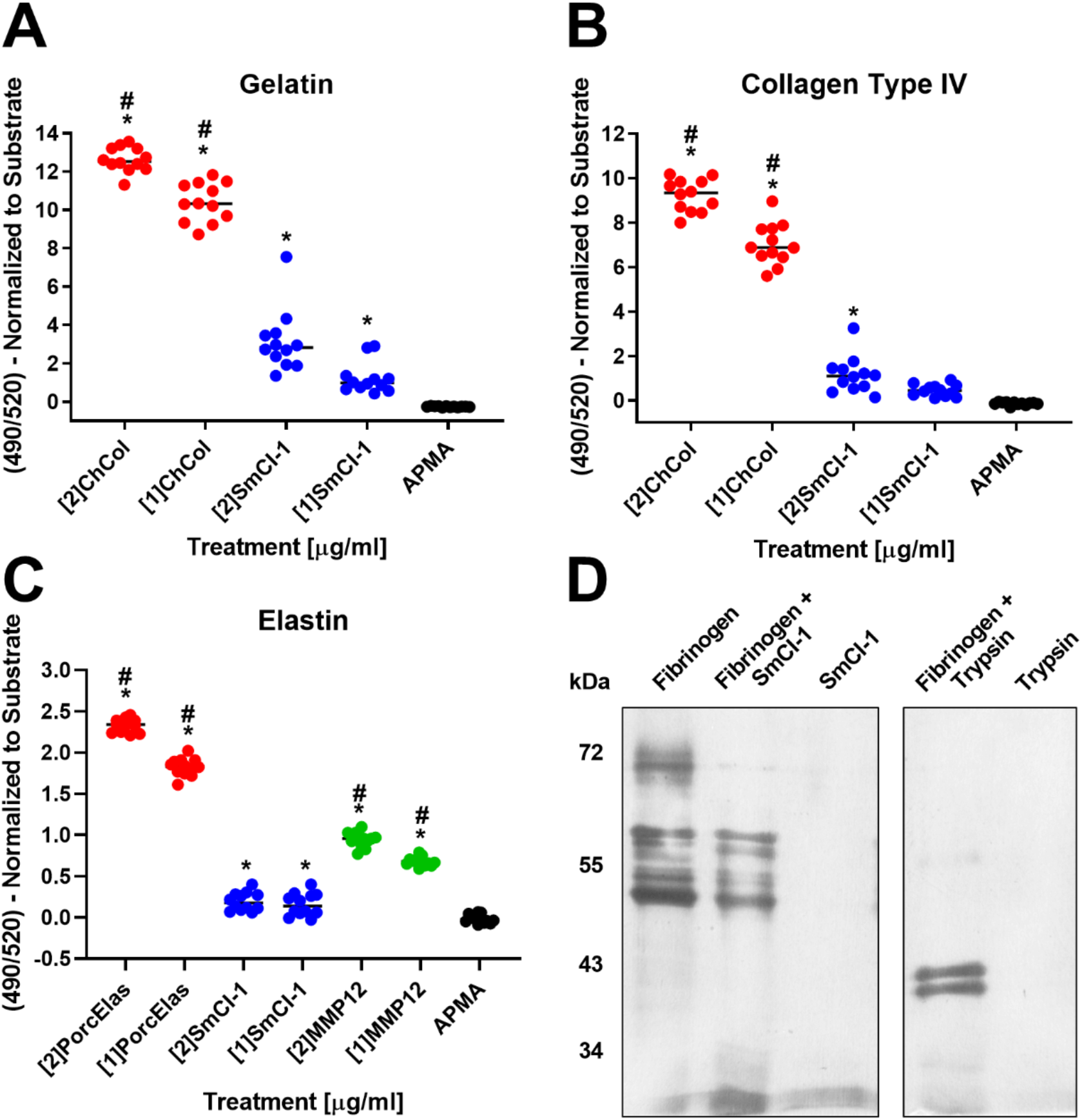
SmCI-1 weakly cleaves key host ECM components. Recombinant SmCI-1 cleaves gelatin (A), collagen type IV (B), and elastin (C). In all cases, known positive controls for collagen type 4 and gelatin cleavage (*C. histolytica* collagenase) and elastin (porcine elastase and human MMP12) displayed higher levels of cleavage. Differences from the APMA vehicle control are denoted using (*) and samples with significantly higher cleavage activity than rSmCI-1 are dented using (#). (D) Silver stain demonstrating the capacity of trypsin to cleave all three subunits of fibrinogen, SmCI-1’s ability to cleave the alpha subunit of fibrinogen only (n=12).

Under reducing SDS-PAGE conditions, fibrinogen separates into the three distinct subunits of which it is composed (α, β, and γ). Our positive control, trypsin cleaves all three subunits, as evidenced by the disappearance of the bands of all three subunits and the appearance of two bands roughly 40kDa in size. Recombinant SmCI-1, alternatively, cleaves only the alpha subunit of fibrinogen, as shown by the disappearance of the α-subunit band at ∼70kDa (Figure 4D).

#### Cleavage of Immunological Substrates

Four immunologically relevant proteins were examined as possible substrates for SmCI-1: complement component C3, CD4, CR1, and Immunoglobulin G (IgG). The later three were all cleaved by trypsin but were unaltered by rSmCI-1 treatment (Figure 5, SFigure 2). The only immunologically-relevant protein examined that was affected by rSmCI-1 treatment was complement component C3b (Figure 5A). Trypsin was capable of cleaving both C3b and the C3a anaphylatoxin.

**Figure 5.**
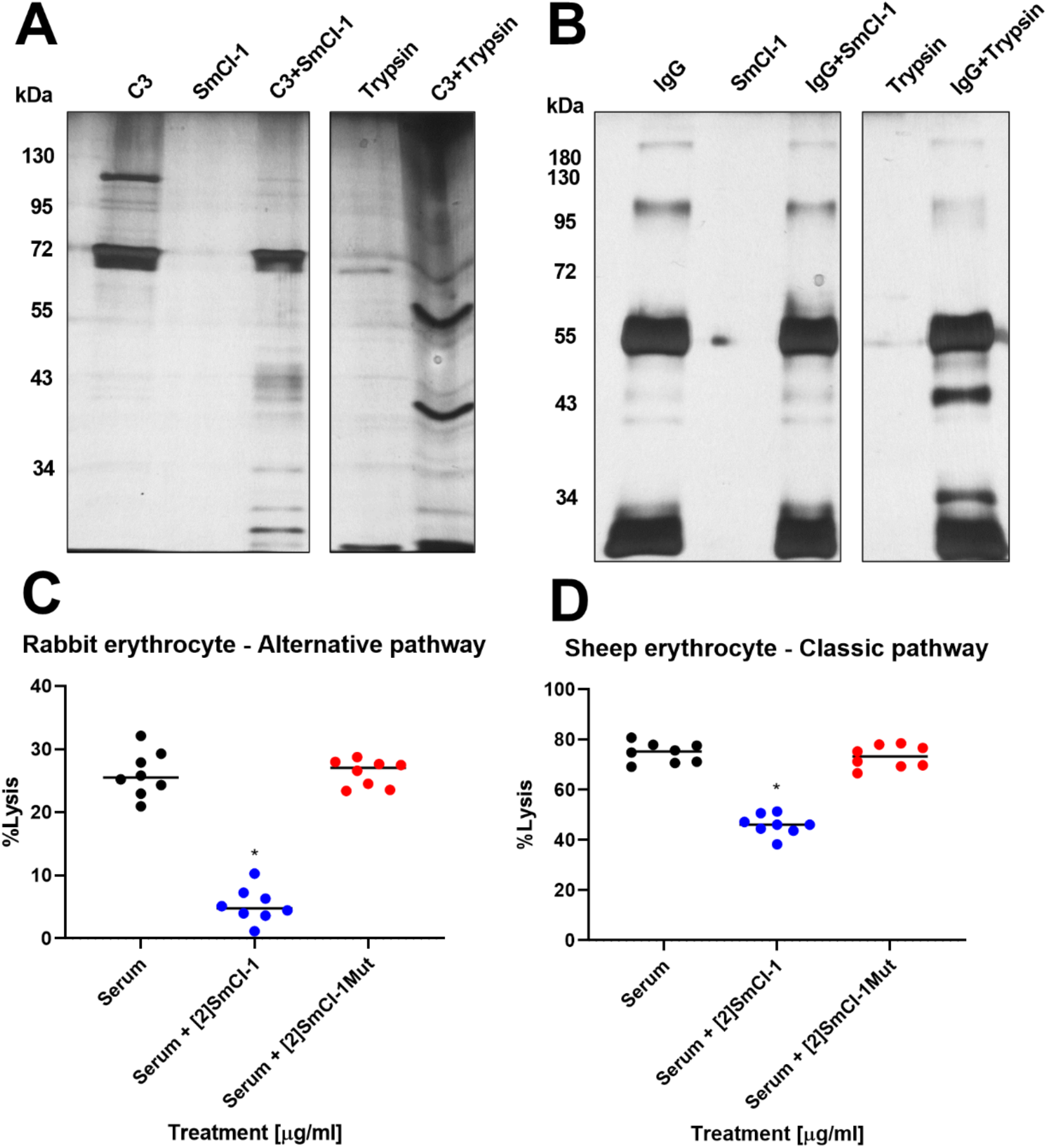
SmCI-1 Cleaves complement component C3. Silver stain of 2μg of (A) human complement component C3 loaded into the lanes of a reducing SDS-Page Gel. Under such conditions, C3 is separated into a C3b band at roughly 110kda, and a C3a band of ∼72kda. Trypsin treatment resulted in the degradation of both C3b and C3a, while SmCI-1 degrades only C3b into fragments visible between 26kda and 55kda. (B) Silver stain of human IgG separated into heavy (∼50kda), and light (∼25kda) chains, with residual complete antibody at higher molecular weights. (C) Rabbit erythrocyte lysis via the alternative complement pathway and (D) Sheep erythrocyte lysis via the classical complement pathway using human serum is decreased by addition of rSmCI-1, but not rMutSmCI-1. Significant differences from untreated cells indicated by (*). (n=8)

The cleavage of complement component C3b led us to examine the effects of SmCI-1 on both the classical and alternative complement pathways (Figure 5C,D). Pre-treatment of human serum with rSmCI-1 caused a significant decrease in alternative pathway-mediated lysis from 26.1±1.3% to 5.3±1.0%. A similar observation was made of the classical pathway, which displayed a significant decrease in lysis from 74.8±1.4% to 46.0±1.4% when treated with rSmCI-1. Treatment with rMutSmCI-1 did not result in a significant decrease in lysis, with 26.2±1.6% and 73.2±0.7% lysis seen in the alternative and classical pathway assays, respectively (Figure 5C,D).

### Cytokine Profiles

SmCI-1 was found to have the capacity to alter the cytokine profiles of human PBMCs, as revealed by cytokine array profiling (STable 1). Amongst this data lie key observations on the effect of rSmCI-1 on the production of key cytokines, as well as whether MMP activity is required for these effects (Figure 7). Whole cercaria lysate (WCL) stimulated cells released 1060±220 pg/ml of Eotaxin-1 after 24 hours. Treatment with rSmCI-1 significantly reduced the amount of Eotaxin-1 to 365±36 pg/ml (p<0.05). In the case of IL-5, WCL stimulated cells produced 330±79 pg/ml, while treatment with rSmCI-1 significantly reduced these levels to 83±16 pg/ml (p<0.01). IL-1β levels were significantly lowered from 355±42 pg/ml to 104±8.8 pg/ml at 24 hours by SmCI-1 addition (p<0.01). Another inflammatory cytokine, IL-12 (quantified in this assay via detection of the IL-12p70 subunit), also saw rSmCI-1 reduce levels from 180±34 pg/ml to 77±13 pg/ml at 24 hours. Eotaxin-1 (1138±301 pg/ml), IL-5 (332±42 pg/ml), IL-1β (394±102 pg/ml), and IL-12 (189±21pg/ml) levels all failed to differ from the WCL only treated cells in a significant manner. Despite lower pg/ml levels overall, the same difference for these four cytokines were also observed at the 12-hour timepoints, with wild-type rSmCI-1 reducing their levels, and the mutant failing to do so.

**Figure 6.**
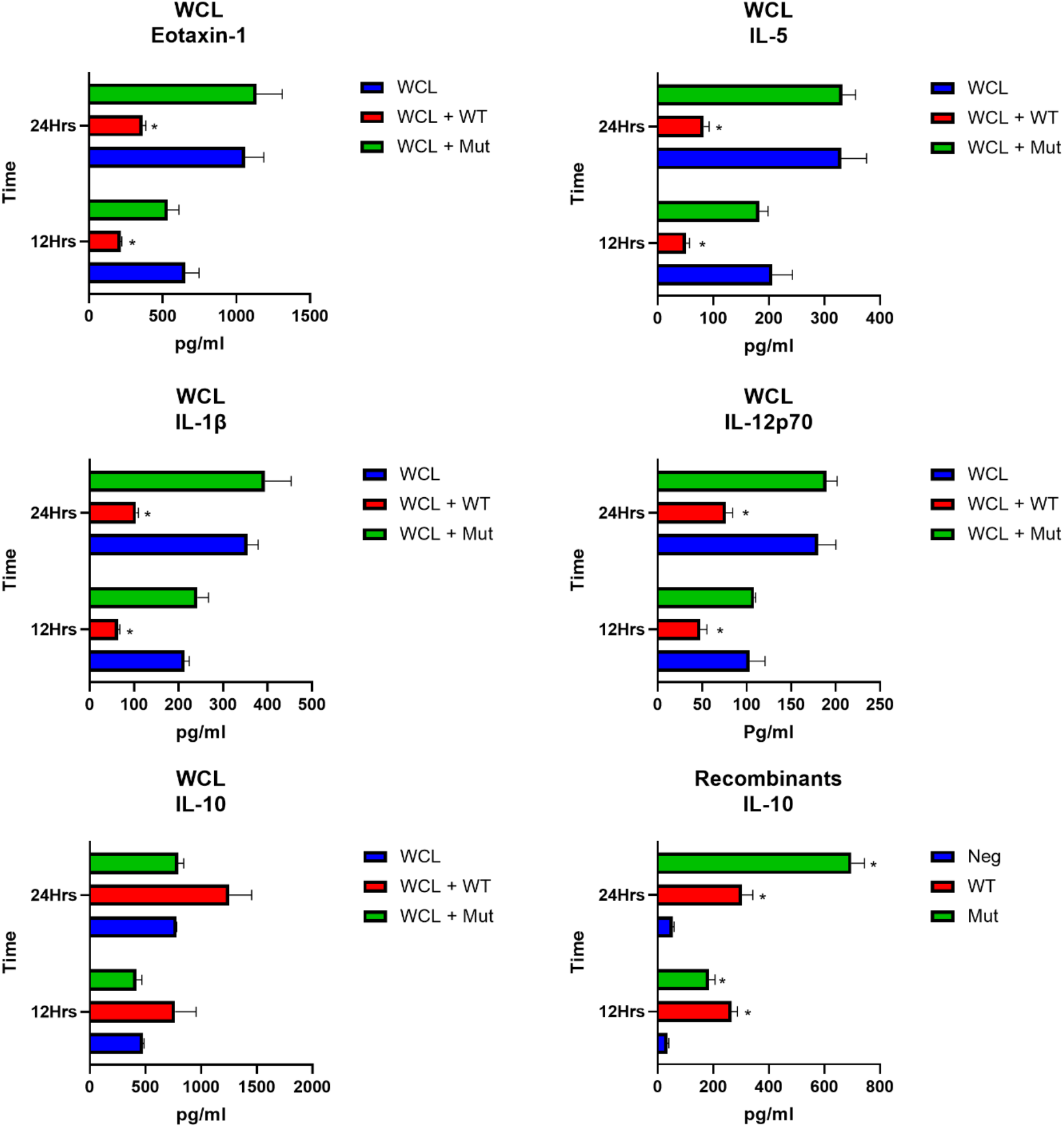
SmCI-1 alters cytokine profiles in an activity dependent manner when stimulated with whole cercarial lysate. Cytokine levels as measured by Luminex panel presented in pg/ml. Differences in levels of key cytokines stimulated with whole cercarial lysate (WCL) totalling the contents of 5 *S. mansoni* cercaria. Treatments with 2μg/ml of either wild type rSmCI-1 (WT) or mutant SmCI-1 (Mut) effects levels of these cytokines. The effect of rSmCI-1 on IL-10 levels features an increase is shown for both cells exposed to WCL, as well as cells exposed only to the recombinants themselves. (*) Indicates significant differences between the treatment and either the WCL control or untreated negative control. (n=3)

**Figure 7.**
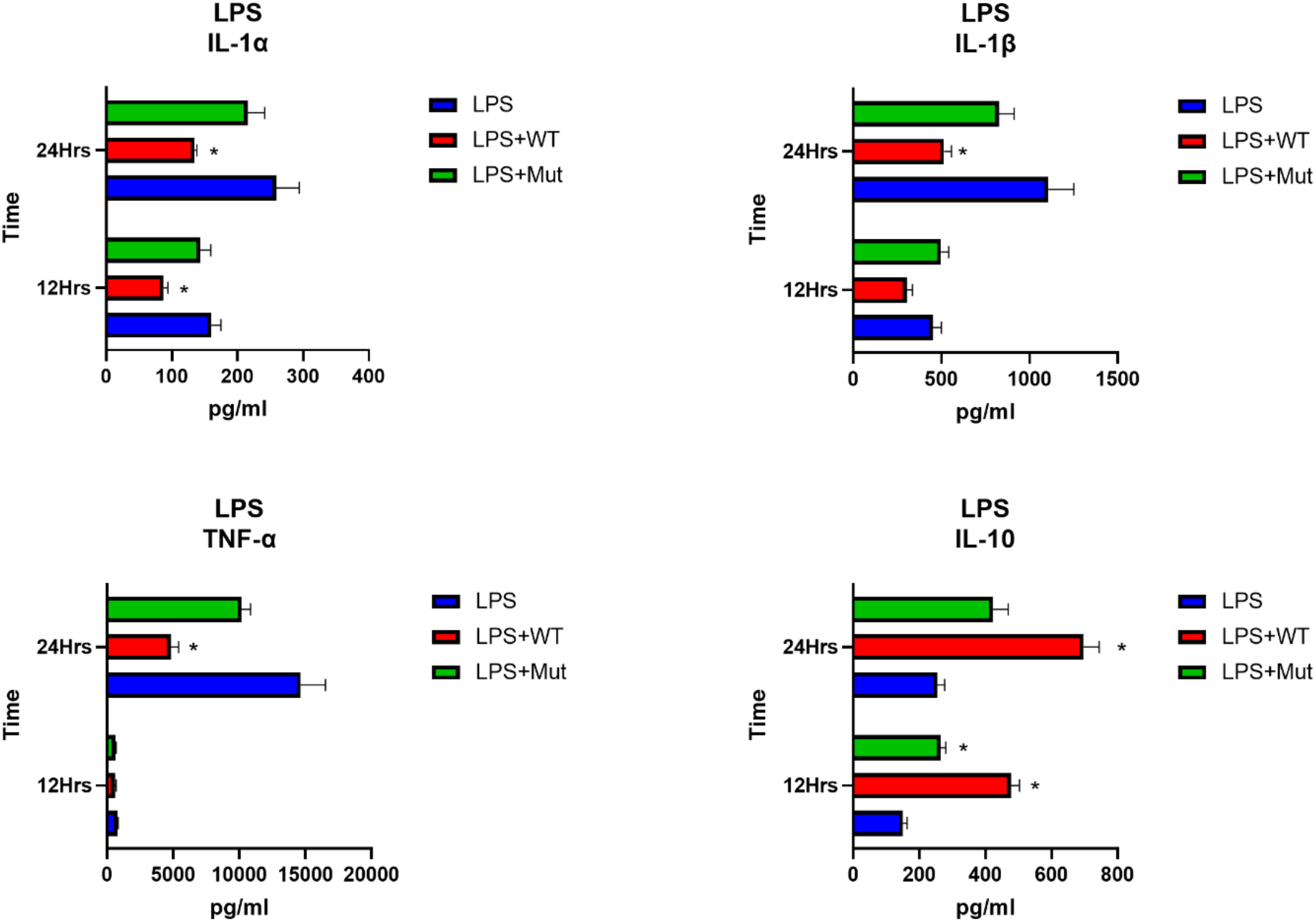
SmCI-1 alters cytokine profiles in an activity dependent manner when stimulated with lipopolysaccharide. Cytokine levels as measured by Luminex panel presented in pg/ml. Differences in levels of key cytokines stimulated with 1μg/ml lipopolysaccharides (LPS). Treatments with 2μg/ml of either wild type rSmCI-1 (WT) or mutant SmCI-1 (Mut) effects levels of these cytokines. (*) Indicates significant differences between the treatment and either the WCL control or untreated negative control. (n=3)

We also examined anti-inflammatory molecules such as IL-10. In the case of IL-10 generated by WCL stimmed PMBCs, our treatments failed to differ significantly from each other, although a general trend towards higher IL-10 production is seen when rSmCI-1 is added (780±4.9 pg/ml vs 1253±348 pg/ml at 12 hours, and 478±17 pg/ml vs 767±329 pg/ml at 24 hours). Regarding PBMCs that were not stimulated with WCLs, but only treated with recombinants, rMutSmCI-1 treatment resulted in the upregulation of numerous cytokines, while rSmCI-1 did not (Stable 1). The only cytokine altered by both rSmCI-1 and rMutSmCI-1 was IL-10, the levels of which were raised at 12 hours from 35±9.0 pg/ml to 266±37 pg/ml and 185±37 pg/ml respectively (P<0.05). The same affects were also seen at 24 hours, with unstimulated levels of IL-10 seen at 54±8.8 pg/ml, increasing significantly to 303±68 pg/ml and 697±82 pg/ml via SmCI-1 and MutSmCI-1 treatment, respectively (p<0.01).

A similar pattern for cytokine alteration was also observed for pro-inflammatory molecules and IL-10 in the context of LPS stimulated PBMCs (Figure 7). IL-1β production was significantly reduced 24hrs post stimulation, with LPS treated cells producing 1107.3±144.5 pg/ml, while cells treated with LPS and rSmCI-1 producing only 515.9±41.5 pg/ml. No significant reduction in IL-1β was seen for LPS+rSmCI-1Mut at 24hrs, or via the addition of either recombinant for the 12hr timepoint. LPS treatment resulted in higher IL-1α and TNF-α levels than those produced by cells exposed to WCL, with observed levels 24hr post LPS stimulation of 259.1±34.7 pg/ml and 14648.4±1878.8 pg/ml. However, IL-1α and TNF-α levels were significantly reduced following addition of rSmCI-1 (p<0.05), but not rSmCI-1Mut. This observation was also observed at 12hr post stimulation. IL-10 increases were measured when adding either recombinant or mutant SmCI-1 to LPS-stimulated cells at both 12 and 24 hour timepoints. These increases were all statistically significant, with the exception of LPS+rSmCI-1Mut at 24 hours post stimulation (p=0.057).

### Cytokine Cleavage

Given the capacity of rSmCI-1, but not rMutSmCI-1, to lower Eotaxin-1 and IL-5 levels, as well as findings in other systems that metalloproteases can cleave Eotaxin-1, we set about examining whether SmCI-1 is able to cleave Eotaxin-1 and IL-5. Neither recombinant cytokine was cleaved by rSmCI-1 (Figure 8). Trypsin completely degraded Eotaxin-1, and partially degraded IL-5.

**Figure 8.**
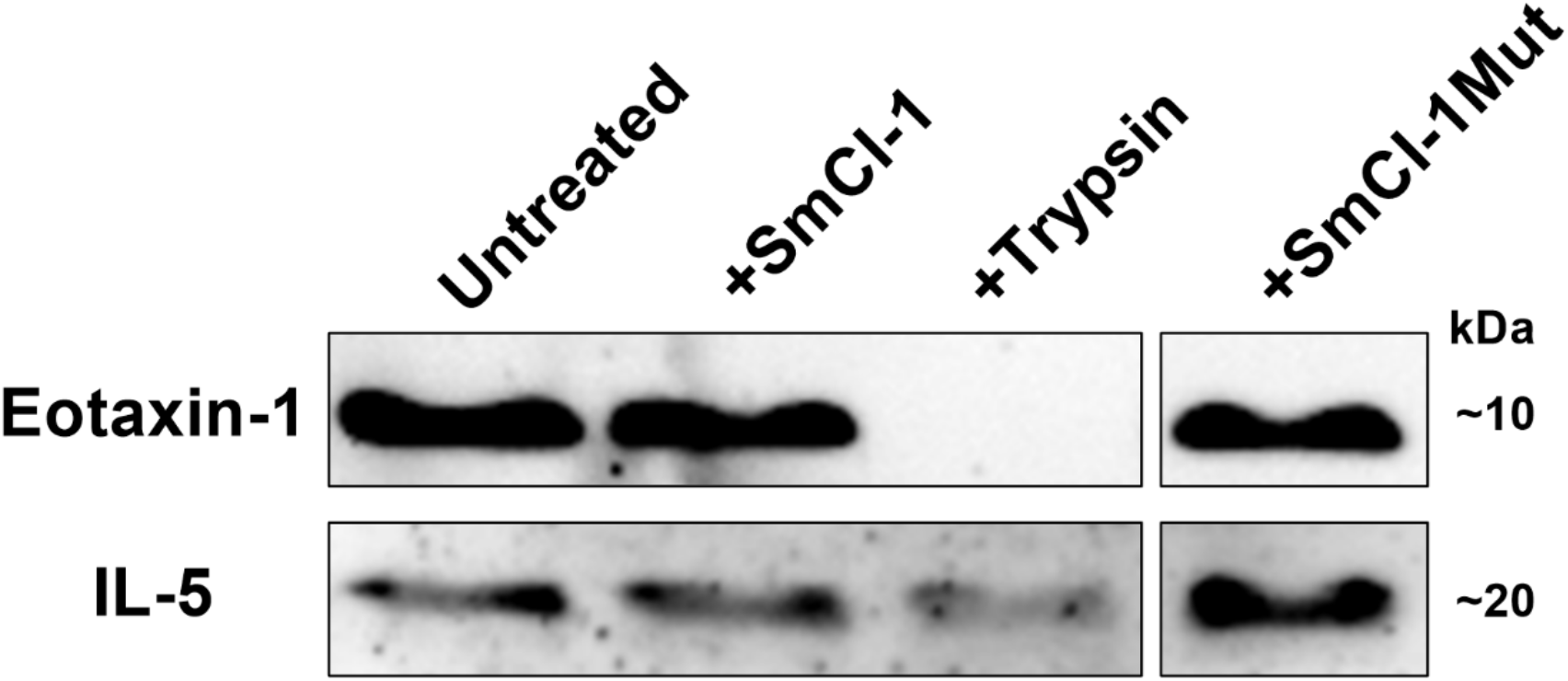
*S. mansoni* cercarial invadolysin-1 do not cleave key eosinophil-associated cytokines. Western blots containing 50ng/lane recombinant human cytokines treated with 1μg/ml of the indicated proteases and probed with relevant monoclonal antibodies. Trypsin degrades Eotaxin-1 while the invadolysins do not. IL-5 was weakly affected by trypsin, and not degraded by either the wild type or mutant rSmCI-1.

**Figure 9.**
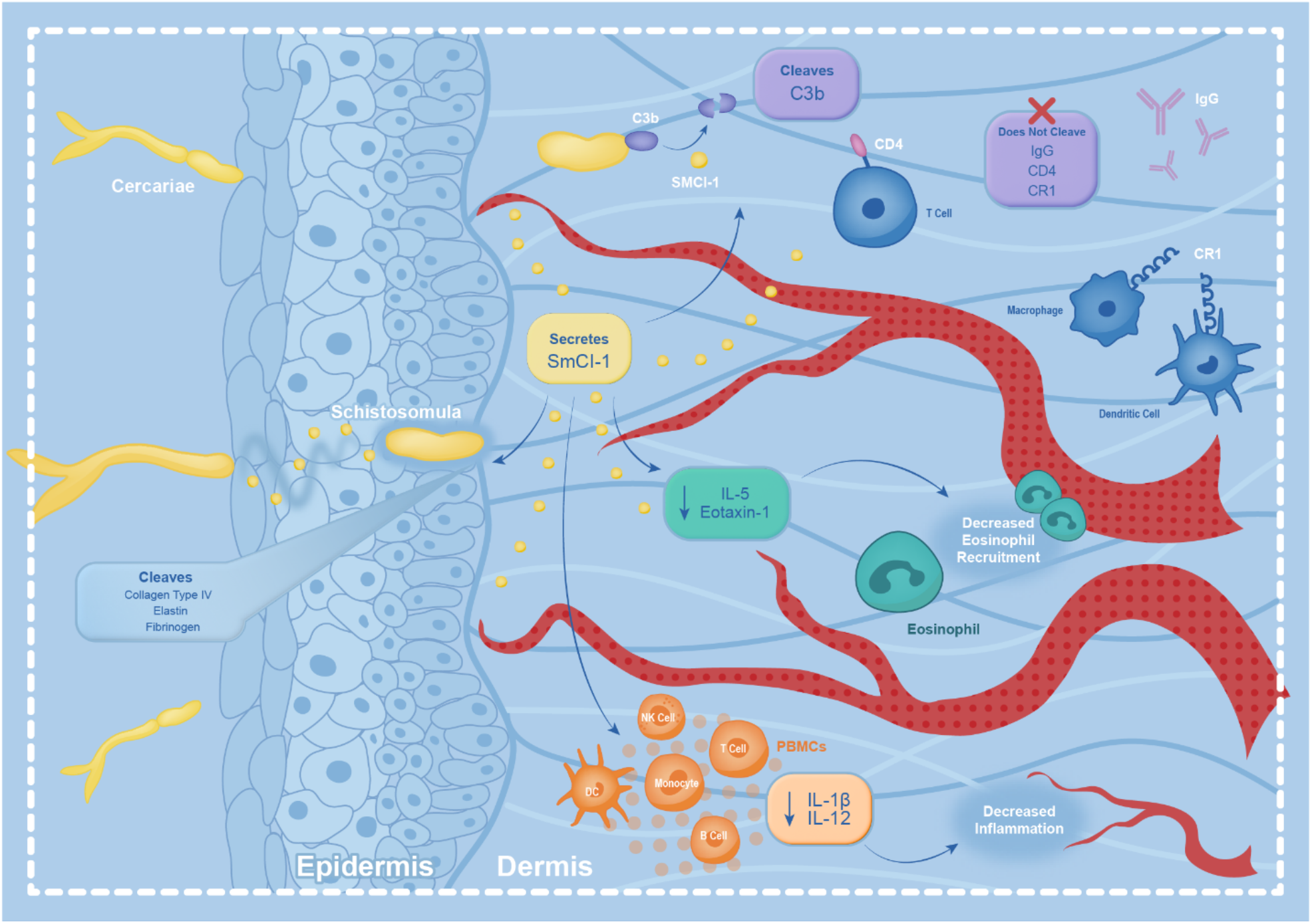
SmCI-1 functions in numerous ways suggesting a key role in host penetration and survival during the initial stages of intra-human infection. These include the degradation of host ECM factors involved in the integrity of the skin. It also includes the targeted cleaving of complement component C3b, which might lead to the reduction of complement mediated lysis of the parasite, as well as reduced opsonization of the larvae, resulting in less immune cell engagement. It also includes the reduction of IL-5 and Eotaxin-1 production, an observation suggesting a role in reducing eosinophil recruitment and development. Finally, the reduction of IL-1β and IL-12 caused by SmCI-1 is indicative of a reduction in inflammation in the skin, which might explain the lack of swelling and site-specific inflammation seen during most *S. mansoni* infections.

## Discussion

Surviving the early stages of infection in human skin, prior to the development of a multi-layered tegument that renders larval schistosomes highly resistant to immune mediated attack, is critical for *S. mansoni* survival (3,59). We sought to characterize the function of the most abundantly produced invadolysin in *S. mansoni* cercaria (13,26). We find compelling evidence that SmCI-1 is a potent immunomodulator that can create an environment conducive to survival of the cercaria during its transformation into a schistosomula.

### The predicted structure of SmCI-1 is most similar to that of GP63

SmCI-1 displays highest amino acid similarity to GP63 from *Leishmania major* to the (57). The predicted three-dimensional structures of both proteins bare a remarkable level of similarity, especially when considering the canonical features of metzincin metalloproteases. Both GP63 and SmCI-1 possess histidine residues that all zinc metalloproteases employ in the coordination of a zinc ion in the active site of the protease (60). Typically, 1-3 amino acids separate the first and second histidine, with a glutamic acid immediately C-terminal to the first histidine (HEXXH) (34,60). The metzincins, a subset of zinc metalloproteases, frequently display an expanded protease motif (HEXXHXXGXXH) with a third histidine employed in coordinating the zinc ion, with some employing as many as 20-120 amino acids between the second and third histidine (34,57,60). While GP63 has 65 amino acids separating its second and third histidines, SmCI-1 has 100. Both possess a methionine predicted to exist underneath the active site, and in the case of SmCI-1, this would be Met347. This methionine is present near to the third histidine in all metzincin proteases, and is the amino acid from which this subset of proteases derives its name (57). The existence of this methionine has been found to be important in stabilizing such proteases, as mutation of this residue can lead to lowered proteolytic activity and increased autolysis (37,61,62). Finally, SmCI-1 has three cysteine residues N-terminal to the active site, as does GP63. While we are not aware which of these residues function in a potential cysteine switch mechanism in either protein, the presence of these amino acids further confirms structural similarities between SmCI-1 and other metzincin invadolysins.

#### SmCI-1 is found in *S. mansoni* acetabular glands and expulsed during transformation

The original identification of SmCI-1 suggests it is a prominent protein of the acetabular gland (26), an observation supported by our immunofluorescence assessment. The placement of SmCI-1 staining inside of the cercaria in a location consistent with the bulk of acetabular contents, as observed by z-stack images, was an expected finding. Our western blotting of cercarial bodies and E/S products also helped to confirm this, as the release of SmCI-1 into RPMI culture media is consistent with the fact that cercaria expulse acetabular gland contents under such conditions (26). We detect the expected ∼65kda protein in Western blot along with a faint second band roughly 49kda in size in cercaria samples. Cleavage of the zymogen into an active form is a near universal requirement for MMPs, thus the ∼49kda likely reflects the presence of an activated form of SmCI-1 retained within the cercaria (38,63,64). SmCI-1 is gradually released from the cercaria between 0- and 4-days post transformation. The exact kinetics of acetabular gland release and the location surrounding the parasite in the skin has long been contested (14,15,19–21). Previous research has demonstrated that acetabular gland contents can be present at the apical end of cercaria as they arrive in the dermis, wherein the are likely to encounter numerous immune cell types (65,66). Additionally, these glands have been observed to atrophy between 48-72 hours post penetration, meaning they are still present as the parasite combats the immune response in the dermis (67). The vortexing transformation method used in this study is not ideal for studying acetabular gland release kinetics given that the cercaria are not squeezed as they would be moving between skin cells. This more natural path likely facilitates a more rapid expulsion of E/S products. However, the more controlled context of culturing suggests that as the parasite releases SmCI-1 over time, it does not seem to continue to produce SmCI-1 to replace that which it has already expulsed. This is consistent with the finding that SmCI-1 is most highly upregulated in the germ ball stage of development, as the cercaria prepares to exit the snail, and not in the emerged cercaria themselves (32).

#### MMP activity of SmCI-1 leads to the cleavage of key host proteins

SmCI-1 clearly displays MMP activity. Whereas activation of MMP-8 (positive control) using APMA significantly increased its activity, the MMP activity of SmCI-1 did not significantly differ between activated and non-activated treatment groups. Possible explanations for this are SmCI-1 being self-activating or activating neighboring SmCI-1 molecules. There is precedent for this line of thought, as *Leishmania sp*. GP63 is capable of cleaving peptides composed of the last 4 amino acids of the protein pro sequence and the first 5 amino acids of the mature GP63, suggesting autoproteolysis as a possible machanism of activation, a finding supported by the fact that zinc chelation and mutation of the HEXXH active site of GP63 reduce its maturation (43). The concept of neighboring invadolysin molecules being responsible for such activation is less likely, given that addition of activated GP63 to GP63 zymogen does not facilitate increased activation of the zymogen (68). The release of SmCI-1 as *S. mansoni* migrates through human skin may benefit from having SmCI-1 not requiring activation, as rapid activity upon expulsion from the acetabular glands would aid in degradation of host components. Alternatively, another possibility could be the activation of SmCI-1 by another acetabular enzyme, although the lack of such enzymes in our studies relying on recombinant proteins suggests that if another enzyme facilitates activation of SmCI-1 during penetration, it is not the only way in which SmCI-1 can be activated. The inhibitory effect of 1,10-phenanthroline further confirmed that SmCI-1 functions as a zinc-based metalloprotease. Finally, mutation of the active site glutamic acid into a non-polar glycine completely abrogated the MMP activity of SmCI-1.

The glutamic acid at this site acts as a nucleophile in activating water molecules, thereby facilitating hydrolysis of substrate proteins (34,69). The mutant SmCI-1 allowed us to further assess the ability of this invadolysin to affect human cells in both an MMP activity dependent and independent manner.

Examination of the proteolysis of several host skin structural molecules by SmCI-1 was predicated upon the knowledge that *S. mansoni* products have been known to degrade host ECM at a distance from the invading larval schistosome (70,71). Of these host molecules, collagen type IV and elastin are of particular relevance, given the abundance of elastin fibres in the human skin dermis that exist as a barrier to invading pathogens, and the large quantity of collagen type IV found in the basement membrane separating the epidermis and dermis, which schistosomes must pass through on their way to finding a venule (72,73). SmCI-1 was capable of cleaving gelatin and collagen type IV. It was, however, less capable of doing so than our positive control, *C. hystolytica* collagenase. The same can be said of its ability to cleave elastin in comparison to porcine elastase and human MMP-12. Given the fact that the serine elastases has been shown to be responsible for the majority of the activity required to enter through human skin, it is possible that the low elastase/collagenase activity demonstrated by SmCI-1 is a function of it not being required for the purpose of degrading these molecules (25). Regarding fibrinogen, while not as abundant in the skin as either collagen or elastin, it may be present at the site of penetration to facilitate wound healing and blood clotting, a function that would likely be negatively impacted by the cleavage of the fibrinogen α-subunit by SmCI-1.

When examining several different immunologically relevant factors in our cleavage assays, we found complement component C3b to be cleaved by SmCI-1. This aligns with studies on GP63 that demonstrate that it can cleave complement component C3. Cleavage of C3 results in an increased capacity for *Leishmania* to survive complement mediated lysis in human serum, which is suggestive of a reason for targeting C3b for cleavage (45,49). *S. mansoni* must also contend with complement-mediated killing by human serum, but only during the period in which they are undergoing shedding of the immunostimulatory glycocalyx (74). The degradation of C3b by SmCI-1 resulted in decreased complement mediated hemolysis via classical and alternative complement pathways. The abundance of SmCI-1 in the acetabular glands, as well as its release during the initial stages of infection, suggests that this invadolysin is likely a key method by which cercaria avoid complement mediated lysis during the early stages of infection. The inability of SmCI-1 to cleave IgG suggests that both the alternative and classical complement pathways were both affected by targeting of C3b, as opposed to targeting multiple humoral opsonins. The likelihood of complement enhanced killing of skin-stage *S. mansoni* is exacerbated by the presence of human eosinophils and neutrophils, which have display schistosome killing activity in vitro, and which is enhanced by the addition of complement factors (75). It is possible that SmCI-1 cleavage of C3b could be a manner by which *S. mansoni* avoids opsonization and subsequent recognition by monocytes/granulocytes during the sensitive skin-penetration period of its lifecycle.

SmCI-1 failed to cleave IgG, CD4, and CR1. We examined IgG, given that schistosomula E/S products, containing metalloproteases, have previously been shown to cleave IgG bound to Fc receptors of *S. mansoni* (76). Serine proteases are a likely alternative IgG-targeting protease to metalloproteases, given that they have already been shown to cleave IgE (77). CD4 investigated because *L. major* GP63 degrades it, and potential CR1 cleavage was assessed because of its known association with C3b (46).

#### The effect of SmCI-1 on cytokine profiles

In response to *S. mansoni* cercarial penetration of the skin, humans produce a variety of immunostimulatory cytokines with the aim of creating an inflammatory milieu in which invading parasites are destroyed. Even keratinocytes participate in this response by releasing IL-1α and IL-1β in response to schistosome penetration (78). Schistosomes respond to this inflammation by releasing numerous anti-immune factors (3). These factor range in form and function, from increasing anti-inflammatory molecules such as IL-10 and reducing production of inflammatory cytokines such as IL-1α and IL-1β, to directly affecting cell function by reducing neutrophil migration, facilitating T cell apoptosis and reducing migration of Langherhan cells to nearby lymph nodes (79–85).

Cytokine profiling following treatment of PBMCs with recombinant SmCI-1 revealed the alteration of numerous cytokines important for an immune response in the skin. Firstly, a reduction in IL-1β and IL-12 was observed following addition of SmCI-1, suggesting a capacity to reduce inflammation at the site of infection. Additionally, we also saw changes in cytokines that have not been previously examined in the skin, with SmCI-1 significantly reducing Eotaxin-1 and IL-5 abundance compared to controls. Given the importance of Eotaxin-1 in recruiting eosinophils to the site of infection, the function of IL-5 to activate eosinophils, and the capacity of eosinophils to kill schistosomulae *in vitro*, the downregulation of these cytokines by SmCI-1 likely assists in avoidance of eosinophil mediated destruction (75,86–88).

Of key interest is the fact that IL-1β, IL-12, Eotaxin-1 and IL-5 were all significantly reduced by the MMP-active form of SmCI-1, but not MutSmCI-1. This lead us to attempt a cleavage assay on two of the cytokines (Eotaxin-1 and IL-5), given the precedent that some hookworm metalloproteases cleave Eotaxin-1 (56). Unlike in hookworms, SmCI-1 failed to cleave eotaxin-1. IL-5 was similarly unaffected. This finding suggests to us that another mechanism may be responsible for the downregulation of these cytokines. Possibilities include the cleavage of surface receptors, the targeting of intracellular signalling molecules or affecting upstream immune stimulating pathways that lead to the production of these cytokines. It is possible that SmCI-1 could cleave cell signalling molecules, given the ability of *L. major* GP63 to cleave MRP and NF-κB, and the recent observations that host macrophages, DCs, and neutrophils are all capable of internalizing schistosome E/S products (65).

In addition to the reduction of proinflammatory cytokines, we also saw an increase in anti-inflammatory cytokines such as IL-10. Significant increases for both wild type and mutant SmCI-1 treated cells were seen at both 12 and 24 hours in unstimulated PBMCs. This suggest that SmCI-1 has the capacity to elicit IL-10 production in a non-MMP activity dependent manner. We did not observe a significant increase in IL-10 post SmCI-1 treatment in WCL stimulated cells, which could reflect the presence of other IL-10-inducing elements overriding the effects of SmCI-1 with respect to IL-10 production, given that levels are significantly higher than what was seen in unstimulated cells. Because our recombinant proteins were produced in HEK cells, it is unlikely that IL-10 production was caused by endotoxin contamination. Despite this fact, we treated PMBCs with pure LPS and did not see Il-10 production as high to that elicited by SmCI-1 treatment (STable1).

Beyond treating PBMCs with recombinant/mutant SmCI-1 or whole cercarial lysate, we also examined the ability of rSmCI-1 to alter the pro-inflammatory state induced by LPS. In addition to reducing IL-1β, as seen with WCL-mediate stimulation, we observed significant reductions in IL-1α and TNF-α when treating LPS-stimulated PBMCs with rSmCI-1. As with WCL treated cells, the measured reduction in the production of these two important pro-inflammatory cytokines was observed for rSmCI-1, but not rSmCI-1Mut. The reduction in these three cytokines suggest that SmCI-1 may be targeting inflammatory pathways common to multiple pathogen types as part of its anti-inflammatory role. The reduction in Il-1α and TNF-α speak strongly to this possibility, given that they were not as upregulated in WCL-stimulated cells as they were in LPS-stimulated cells, suggesting SmCI-1 can downregulate pro-inflammatory responses that are not strongly upregulated by the schistosome cercaria itself. Understanding the anti-inflammatory properties of helminth produced factors is a growing area of research that is continuously expanding, building upon the hygiene/old friends hypothesis in an attempt to ultimately utilize helminth-derived molecules as anti-inflammatory therapeutics for the treatment of inflammatory diseases (89–92). The ability of SmCI-1 to reduce LPS-mediated inflammation positions it within this growing group of anti-inflammatory factors. While different receptors and signalling pathways could be suggested as possible targets for SmCI-1 interference, the identification of such remains to be undertaken, and will be a significant focus of our research moving forwards.

Interpretation of the cytokine array results require several caveats. First off, while PBMCs do have numerous cell types capable of giving us valuable information, they lack keratinocytes, macrophages, and granulocytes, so the possibility exists that we are not capturing the entirety of the effects of SmCI-1 observed *in vivo*. Secondly, these assays are performed *in vitro*, with more cells than are expected to encounter an invading schistosome during a natural infection, resulting in higher pg/ml levels than might be expected in the context of migration through human skin. Given that other research into the effects of *S. mansoni* immunomodulators found in the skin have employed mouse cells, as well as keratinocyte cell lines, we felt that the use of human PBMCs would allow for a more host-relevant understanding and wider capture of the effects of SmCI-1 on a variety of combined cell types (78,80–82). This reduction of inflammatory factors is an intriguing observation given the observation that *S. mansoni* cercaria often do not elicit substantial cecarial dermatitis, unlike exposures to other mammalian and avian schistosomes (93). We propose that some host specificity of penetration gland proteases such as invadolysins may play a role in this difference in inflammation.

## Conclusion

SmCI-1 is the second most abundant acetabular gland protein released by *S. mansoni* cercaria during penetration into human skin. This fact alone merits an examination of its roles during infection. Here, we show that SmCI-1 possesses the hallmark structure and sequences of an invadolysin, a family of proteases shown to be important for infection success in numerous parasites. We demonstrate that SmCI-1 has numerous functions relevant to survival in human skin, including cleavage of structural components, targeting of complement component C3b for cleavage, and subsequent reduction of complement pathway activity. We also show that it can alter the immunological environment in human cells by reducing IL-1β, IL-12, Il-5, and Eotaxin-1 levels, but only while in possession of MMP activity. Additionally, it can increase IL-10 levels, and MMP activity is not necessary for this function. These functions suggest SmCI-1 as an important factor for parasite survival early on during infection (Figure 8). Future work should seek to determine the method by which SmCI-1 enacts its effect on cytokines and seek to determine whether or not it is essential for survival *in vivo*.

## Acknowledgements

Microscopy was performed in the University of Alberta, Faculty of Medicine and Dentistry Cell Imaging Core

## Supporting Information

**Supplementary Figure 1.**
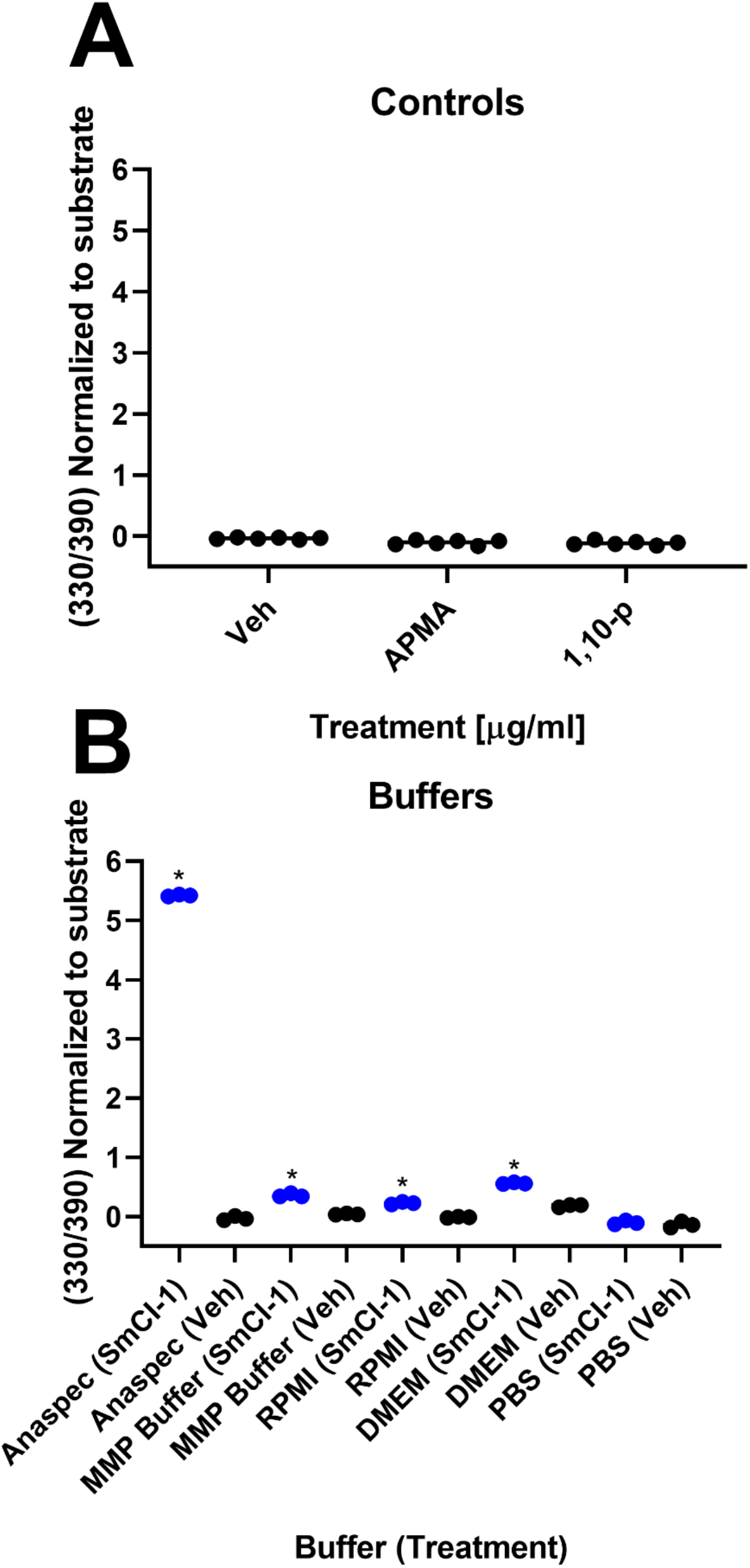
SmCI-1 is active in various buffers. Vehicle, APMA, and 1,10-phenanthroline controls for generic MMP assay fail to vary significantly from substrate controls (A). SmCI-1 activity as measured using a generic fluorometric MMP assay in a variety of buffers (B). The highest activity is seen in the Sensolyte MMP assay provided buffer. Activity is also seen in our lab made generic MMP buffer (50mM Tris, 10mM CaCl_2,_ 150mM NaCl, pH 7.5), as well as RPMI and DMEM. Activity is not seen in Krebs-Ringer Phosphate Buffer (KRPG) (145mM NaCl, 6mM Na_3_PO_4_, 5mM KCl, 0.5mM CaCl_2_, 1mM MgSO_4,_ pH 7.4) or PBS. Statistically significant differences from vehicle only controls indicated using (*).

**Supplementary Figure 2.**
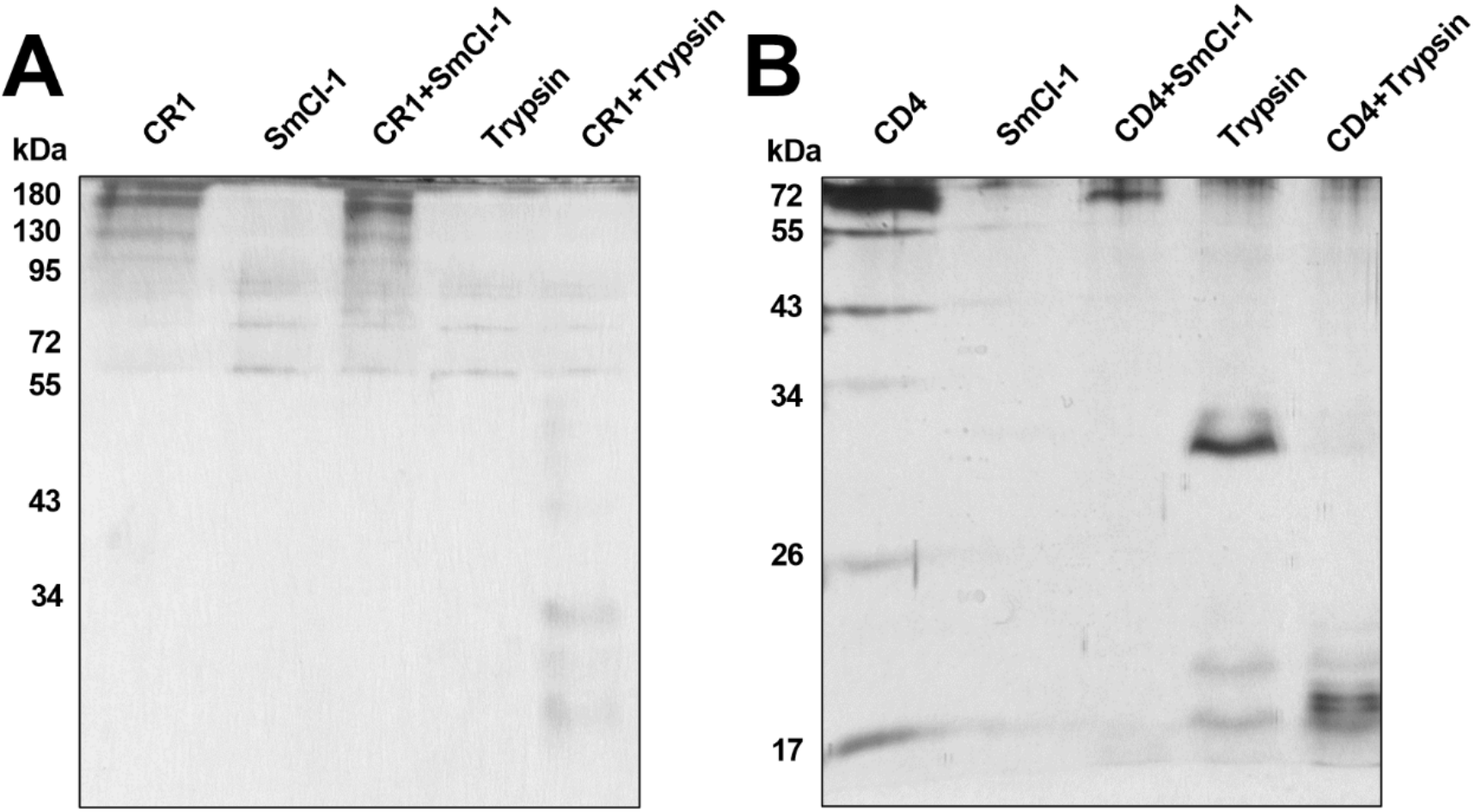
SmCI-1. SmCI-1 fails to cleave CR1 (A), or CD4 (B). Trypsin does cleave these molecules, as evidenced by the appearance of novel bands such as the numerous bands located between 26 and 24kda when it is added to CR1, and the ∼20kda bands seen when trypsin is added to CD4.

**Supplementary Table 1.**
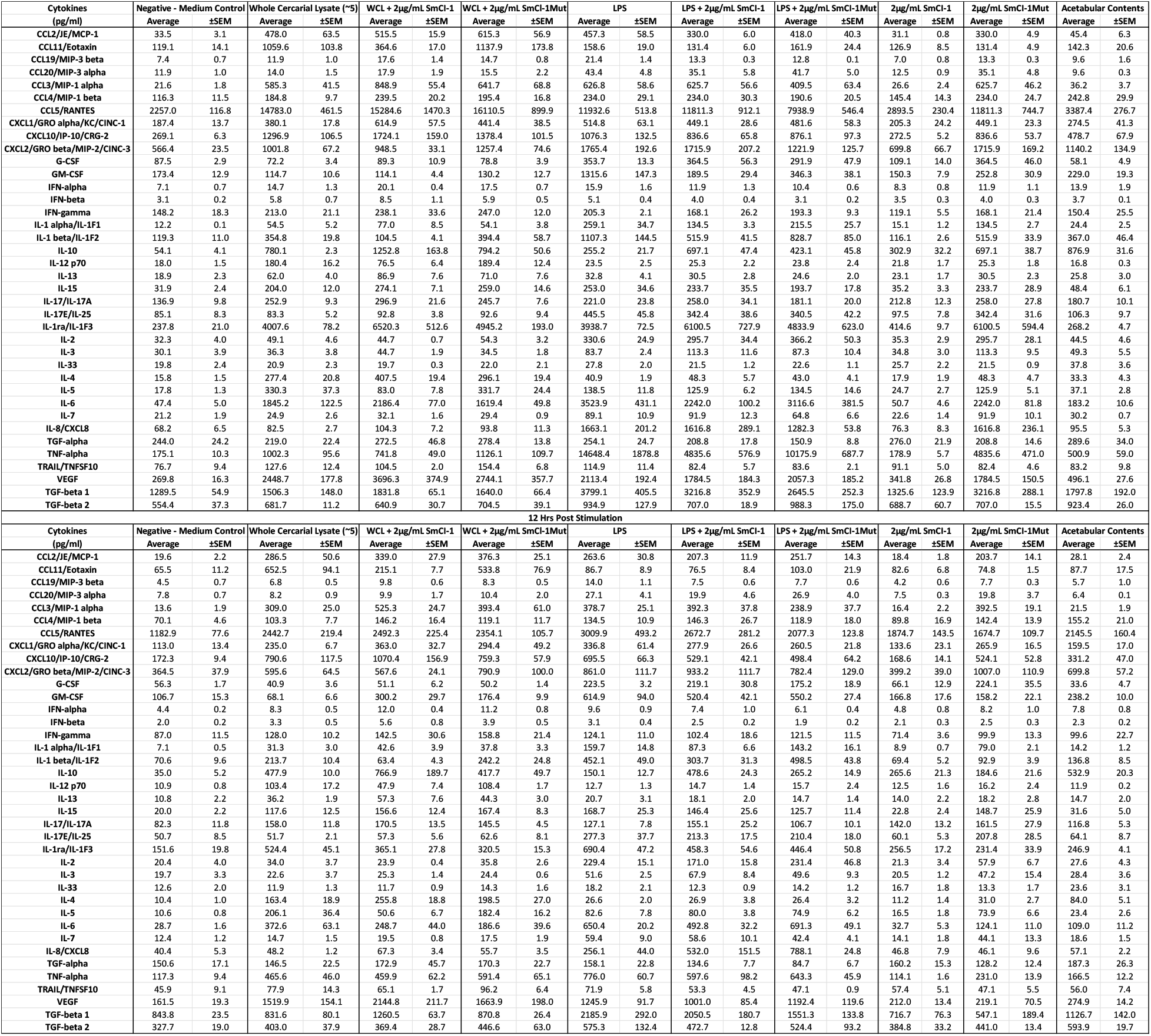
Cytokine array data for PBMCs. Average pg/ml concentrations of individually indicated cytokines treated with *S. mansoni* whole cercarial lysate, acetabular gland contents), LPS, recombinant SmCI-1, or recombinant MutSmCI-1. Values are indicated for both 24 and 12 hour timepoints. Accompanied by Standard error of the mean values (SEM). n=3

## Works cited

1. World Health Organization. The Weekly Epidemiological Record (WER) (2018). WHO. (45):445–52.

2. Pila EA, Li H, Hambrook JR, Wu X, Hanington PC. Schistosomiasis from a Snail’s Perspective: Advances in Snail Immunity (2017). Trends Parasitol. 33(11):845–57. DOI:10.1016/j.pt.2017.07.006

3. Hambrook JR, Hanington PC. Immune Evasion Strategies of Schistosomes (2021). Front Immunol. 11(1):3820. DOI:10.3389/fimmu.2020.624178

4. Angeles JMM, Mercado VJP, Rivera PT. Behind Enemy Lines: Immunomodulatory Armamentarium of the Schistosome Parasite (2020). Vol. 11, Frontiers in Immunology. Frontiers Media S.A.; 2020. p. 1018. DOI:10.3389/fimmu.2020.01018

5. Mitchell KM, Mutapi F, Savill NJ, Woolhouse MEJ. Protective immunity to Schistosoma haematobium infection is primarily an anti-fecundity response stimulated by the death of adult worms (2012). Proc Natl Acad Sci U S A. 109(33):13347–52. DOI:10.1073/pnas.1121051109

6. Butterworth AE. Immunological aspects of human schistosomiasis [Internet] (1998). Vol. 54, British Medical Bulletin. Oxford Academic; 1998 [cited 2022 Feb 1]. p. 357–68. DOI:10.1093/oxfordjournals.bmb.a011693

7. Amaral MS, Santos DW, Pereira ASA, Tahira AC, Malvezzi JVM, Miyasato PA, et al. Rhesus macaques self-curing from a schistosome infection can display complete immunity to challenge (2021). Nat Commun. 12(1):1–17. DOI:10.1038/s41467-021-26497-0

8. Houlder EL, Costain AH, Cook PC, MacDonald AS. Schistosomes in the Lung: Immunobiology and Opportunity (2021). Vol. 12, Frontiers in Immunology. Frontiers Media S.A.; 2021. p. 1330. DOI:10.3389/fimmu.2021.635513

9. McKerrow JH, Doenhoff MJ. Schistosome proteases (1988). Parasitol Today. 4(12):334–40. DOI:10.1016/0169-4758(88)90002-6

10. Vermeire JJ, Taft AS, Hoffmann KF, Fitzpatrick JM, Yoshino TP. Schistosoma mansoni: DNA microarray gene expression profiling during the miracidium-to-mother sporocyst transformation (2006). Mol Biochem Parasitol. 147(1):39–47. DOI:10.1016/j.molbiopara.2006.01.006

11. Taft AS, Vermeire JJ, Bernier J, Birkeland SR, Cipriano MJ, Papa AR, et al. Transcriptome analysis of Schistosoma mansoni larval development using serial analysis of gene expression (SAGE) (2009). Parasitology. 136(5):469–85. DOI:10.1017/S0031182009005733

12. Wu X-J, Sabat G, Brown JF, Zhang M, Taft A, Peterson N, et al. Proteomic analysis of Schistosoma mansoni proteins released during in vitro miracidium-to-sporocyst transformation (2009). Mol Biochem Parasitol. 164:32–44. DOI:10.1016/j.molbiopara.2008.11.005

13. Hambrook JR, Kaboré AL, Pila EA, Hanington PC. A metalloprotease produced by larval Schistosoma mansoni facilitates infection establishment and maintenance in the snail host by interfering with immune cell function (2018). Jolly ER, editor. PLoS Pathog. 14(10):e1007393. DOI:10.1371/journal.ppat.1007393

14. Bartlett A, Brown M, Marriott C, Whitfield” PJ. The infection of human skin by schistosome cercariae : studies using Franz cells (2000). Parasitology. 1121:49–54.

15. McKerrow JH, Salter J. Invasion of skin by Schistosoma cercariae (2002). Trends Parasitol. 18(5):193–5. DOI:10.1016/S1471-4922(02)02309-7

16. Wang Q, Da’dara AA, Skelly PJ. The human blood parasite Schistosoma mansoni expresses extracellular tegumental calpains that cleave the blood clotting protein fibronectin (2017). Sci Rep. 7(1):1–13. DOI:10.1038/s41598-017-13141-5

17. Da’dara AA, Bhardwaj R, Ali YBM, Skelly PJ. Schistosome tegumental ecto-apyrase (SmATPDase1) degrades exogenous pro-inflammatory and pro-thrombotic nucleotides (2014). PeerJ. 2014(1):e316. DOI:10.7717/peerj.316

18. Leontovyč A, Ulrychová L, O’Donoghue AJ, Vondrášek J, Marešová L, Hubálek M, et al. SmSP2: A serine protease secreted by the blood fluke pathogen Schistosoma mansoni with anti-hemostatic properties (2018). Greenberg RM, editor. PLoS Negl Trop Dis. 12(4):e0006446. DOI:10.1371/journal.pntd.0006446

19. Curwen RS, Wilson RA. Invasion of skin by schistosome cercariae: Some neglected facts (2003). Trends Parasitol. 19(2):63–6. DOI:10.1016/S1471-4922(02)00019-3

20. McKerrow JJ. Invasion of skin by schistosome cercariae: Some neglected facts - Response from James J. McKerrow (2003). Trends Parasitol. 19(2):66–8. DOI:10.1016/S1471-4922(02)00018-1

21. Whitfield PJ, Bartlett A, Brown MB, Marriott C. Invasion by schistosome cercariae: Studies with human skin explants (2003). Trends Parasitol. 19(8):339–40. DOI:10.1016/S1471-4922(03)00143-0

22. Marikovsky M, Fishelson Z, Arnon R. Purification and characterization of proteases secreted by transforming schistosomula of Schistosoma mansoni (1988). Mol Biochem Parasitol. 30(1):45–54. DOI:10.1016/0166-6851(88)90131-4

23. Chavez-Olortegu C, Tavares CAP, Resende M. Purification and characterization of a 47 kDa protease from Schistosoma mansoni cerearial secretion (1992). Parasitology. 105(2):211–8. DOI:10.1017/S0031182000074138

24. Newport GR, McKerrow JH, Hedstrom R, Petitt M, McGarrigle L, Barr PJ, et al. Cloning of the proteinase that facilitates infection by schistosome parasites (1988). J Biol Chem. 263(26):13179–84.

25. Salter JP, Lim KC, Hansell E, Hsieh I, McKerrow JH. Schistosome invasion of human skin and degradation of dermal elastin are mediated by a single serine protease (2000). J Biol Chem. 275(49):38667–73. DOI:10.1074/jbc.M006997200

26. Curwen RS, Ashton PD, Sundaralingam S, Wilson RA. Identification of Novel Proteases and Immunomodulators in the Secretions of Schistosome Cercariae That Facilitate Host Entry (2006). Mol Cell Proteomics. 5(5):835–44. DOI:10.1074/mcp.M500313-MCP200

27. Lim KC, Sun E, Bahgat M, Bucks D, Richard G, Hinz RS, et al. Blockage of skin invasion by schistosome cercariae by serine protease inhibitors (1999). Am J Trop Med Hyg. 60(3):487–92. DOI:10.4269/ajtmh.1999.60.487

28. Dalton JP, Clough KA, Jones MK, Brindley PJ. The cysteine proteinases of Schistosoma mansoni cercariae (1997). Parasitology. 114(2):105–12. DOI:10.1017/S003118209600830X

29. Dvořák J, Mashiyama ST, Braschi S, Sajid M, Knudsen GM, Hansell E, et al. Differential use of protease families for invasion by schistosome cercariae (2008). Biochimie. 90(2):345–58. DOI:10.1016/j.biochi.2007.08.013

30. Verjovski-Almeida S, DeMarco R, Martins EAL, Guimarães PEM, Ojopi EPB, Paquola ACM, et al. Transcriptome analysis of the acoelomate human parasite Schistosoma mansoni (2003). Nat Genet. 35(2):148–57. DOI:10.1038/ng1237

31. Berriman M, Haas BJ, Loverde PT, Wilson RA, Dillon GP, Cerqueira GC, et al. The genome of the blood fluke Schistosoma mansoni (2009). Nature. 460(7253):352–8. DOI:10.1038/nature08160

32. Parker-Manuel SJ, Ivens AC, Dillon GP, Wilson RA. Gene expression patterns in larval Schistosoma mansoni associated with infection of the mammalian host (2011). PLoS Negl Trop Dis. 5(8):e1274. DOI:10.1371/journal.pntd.0001274

33. Nagase H. Metalloproteases (2013). Encycl Biol Chem Second Ed. :86–9. DOI:10.1016/B978-0-12-378630-2.00018-9

34. Vallee BL, Auld DS. Active-site zinc ligands and activated H2O of zinc enzymes (1990). Proc Natl Acad Sci U S A. 87(1):220–4. DOI:10.1073/pnas.87.1.220

35. Van Wart HE, Birkedal-Hansen H. The cysteine switch: A principle of regulation of metalloproteinase activity with potential applicability to the entire matrix metalloproteinase gene family (1990). Proc Natl Acad Sci U S A. 87(14):5578–82. DOI:10.1073/pnas.87.14.5578

36. Klein T, Bischoff R. Physiology and pathophysiology of matrix metalloproteases (2011). Amino Acids. 41(2):271–90. DOI:10.1007/s00726-010-0689-x

37. Tallant C, García-Castellanos R, Baumann U, Gomis-Rüth FX. On the relevance of the met-turn methionine in metzincins (2010). J Biol Chem. 285(18):13951–7. DOI:10.1074/jbc.M109.083378

38. Gomiz-Rüth FX. Catalytic domain architecture of metzincin metalloproteases (2009). J Biol Chem. 284(23):15353–7. DOI:10.1074/jbc.R800069200

39. Liu S, Cai P, Piao X, Hou N, Zhou X, Wu C, et al. Expression Profile of the Schistosoma japonicum Degradome Reveals Differential Protease Expression Patterns and Potential Anti-schistosomal Intervention Targets (2014). PLoS Comput Biol. 10(10):e1003856. DOI:10.1371/journal.pcbi.1003856

40. Bouvier J, Schneider P, Etges R. Leishmanolysin: Surface metalloproteinase of Leishmania (1995). Methods Enzymol. 248:614–33. DOI:10.1016/0076-6879(95)48039-0

41. Etges R, Bouvier J, Bordier C. The major surface protein of Leishmania promastigotes is a protease (1986). J Biol Chem. 261(20):9098–101. DOI:10.1016/S0021-9258(18)67621-5

42. McGwire BS, Chang KP, Engman DM. Migration through the extracellular matrix by the parasitic protozoan Leishmania is enhanced by surface metalloprotease gp63 (2003). Infect Immun. 71(2):1008–10. DOI:10.1128/IAI.71.2.1008-1010.2003

43. Bouvier J, Schneider P, Etges R, Bordier C. Peptide Substrate Specificity of the Membrane-Bound Metalloprotease of Leishmania (1990). Biochemistry. 29:10113–9.

44. Brittingham A, Morrison CJ, McMaster WR, McGwire BS, Chang KP, Mosser DM. Role of the Leishmania surface protease gp63 in complement fixation, cell adhesion, and resistance to complement-mediated lysis. (1995). J Immunol. 155(6):3102–11.

45. Joshi PB, Kelly BL, Kamhawi S, Sacks DL, Mcmaster WR. Targeted gene deletion in Leishmania major identifies leishmanolysin (GP63) as a virulence factor (2002). Mol Biochem Parasitol. 120:33–40.

46. Hey AS, Theander TG, Hviid L, Hazrati SM, Kemp M, Kharazmi A. The major surface glycoprotein (gp63) from Leishmania major and Leishmania donovani cleaves CD4 molecules on human T cells. (1994). J Immunol. 152(9):4542–8.

47. Yao C, Donelson JE, Wilson ME. The major surface protease (MSP or GP63) of Leishmania sp. Biosynthesis, regulation of expression, and function (2003). Mol Biochem Parasitol. 132:1–16. DOI:10.1016/S0166-6851(03)00211-1

48. Mosser DM, Edelson PJ. The mouse macrophage receptor for C3bi (CR3) is a major mechanism in the phagocytosis of Leishmania promastigotes. (1985). J Immunol. 135(4):2785–9.

49. Chaudhuri G, Chang K-P. Acid protease activity of a major surface membrane glycoprotein (gp63) from Leishmania mexicana promastigotes [Internet] (1988). Vol. 27, Molecular and Biochemical Parasitology. 1988 [cited 2019 Mar 19].

50. Chaudhuri G, Chaudhuri M, Pan A, Chang KP. Surface acid proteinase (gp63) of Leishmania mexicana. A metalloenzyme capable of protecting liposome-encapsulated proteins from phagolysosomal degradation by macrophages (1989). J Biol Chem. 264(13):7483–9. DOI:10.1016/S0021-9258(18)83260-4

51. Corradin S, Ransijn A, Corradin G, Roggero MA, Schmitz AAP, Schneider P, et al. MARCKS-related protein (MRP) is a substrate for the Leishmania major surface protease leishmanolysin (gp63) (1999). J Biol Chem. 274(36):25411–8. DOI:10.1074/jbc.274.36.25411

52. Gregory DJ, Godbout M, Contreras I, Forget G, Olivier M. A novel form of NF-κB is induced by Leishmania infection: Involvement in macrophage gene expression (2008). Eur J Immunol. 38(4):1071–81. DOI:10.1002/eji.200737586

53. Ahmed AA, Wahbi A, Nordlind K, Kharazmi A, Sundqvist KG, Mutt V, et al. In vitro Leishmania major promastigote-induced macrophage migration is modulated by sensory and autonomic neuropeptides (1998). Scand J Immunol. 48(1):79–85. DOI:10.1046/j.1365-3083.1998.00380.x

54. SØRensen AL, Hey AS, Kharazmi A. Leishmania major surface protease Gp63 interferes with the function of human monocytes and neutrophils in vitro (1994). APMIS. 102(1– 6):265–71. DOI:10.1111/j.1699-0463.1994.tb04874.x

55. Lieke T, Nylén S, Eidsmo L, McMaster WR, Mohammadi AM, Khamesipour A, et al. Leishmania surface protein gp63 binds directly to human natural killer cells and inhibits proliferation (2008). Clin Exp Immunol. 153(2):221–30. DOI:10.1111/j.1365-2249.2008.03687.x

56. Culley FJ, Brown A, Conroy DM, Sabroe I, Pritchard DI, Williams TJ. Eotaxin Is Specifically Cleaved by Hookworm Metalloproteases Preventing Its Action In Vitro and In Vivo (2000). J Immunol. 165(11):6447–53. DOI:10.4049/jimmunol.165.11.6447

57. Schlagenhauf E, Etges R, Metcalf P. The crystal structure of the Leishmania major surface proteinase leishmanolysin (gp63) (1998). Structure. 6(8):1035–46. DOI:10.1016/S0969-2126(98)00104-X

58. Morgan BP. Complement Methods and Protocols [Internet] (2000). Complement Methods and Protocols. 2000 [cited 2022 Aug 18]. DOI:10.1385/159259056x

59. Hockley DJ, McLaren DJ. Schistosoma mansoni: Changes in the outer membrane of the tegument during development from cercaria to adult worm (1973). Int J Parasitol. 3(1):13–20. DOI:10.1016/0020-7519(73)90004-0

60. Rawlings ND, Barrett AJ. Evolutionary families of metallopeptidases (1995). Methods Enzymol. 248:183–228. DOI:10.1016/0076-6879(95)48015-3

61. Butler GS, Tam EM, Overall CM. The canonical methionine 392 of matrix metalloproteinase 2 (Gelatinase A) is not required for catalytic efficiency or structural integrity: Probing the role of the methionine-turn in the metzincin metalloprotease superfamily (2004). J Biol Chem. 279(15):15615–20. DOI:10.1074/jbc.M312727200

62. Pieper M, Betz M, Budisa N, Gomis-Rüth FX, Bode W, Tschesche H. Expression, purification, characterization, and X-ray analysis of selenomethionine 215 variant of leukocyte collagenase (1997). J Protein Chem. 16(6):637–50. DOI:10.1023/A:1026327125333

63. Massova I, Kotra LP, Fridman R, Mobashery S. Matrix metalloproteinases: structures, evolution, and diversification (1998). FASEB J. 12(12):1075–95.

64. K.A. H, T.F. P, G.I. G, J.P. T, D.G. S, R.M. S, et al. Human neutrophil collagenase. A distinct gene product with homology to other matrix metalloproteinases (1990). J Biol Chem. 265(20):11421–4.

65. Paveley RA, Aynsley SA, Cook PC, Turner JD, Mountford AP. Fluorescent imaging of antigen released by a skin-invading helminth reveals differential uptake and activation profiles by antigen presenting cells (2009). Jones MK, editor. PLoS Negl Trop Dis. 3(10):e528. DOI:10.1371/journal.pntd.0000528

66. Fishelson Z, Amiri P, Friend DS, Marikovsky M, Petitt M, Newport G, et al. Schistosoma mansoni: Cell-specific expression and secretion of a serine protease during development of cercariae (1992). Exp Parasitol. 75(1):87–98. DOI:10.1016/0014-4894(92)90124-S

67. Crabtree JE, Wilson RA. Schistosoma Mansoni: An Ultrastructural Examination Of Pulmonary Migration (1986). Parasitology. 92(2):343–54. DOI:10.1017/S0031182000064118

68. Macdonald MH, Morrison CJ, McMaster WR. Analysis of the active site and activation mechanism of the Leishmania surface metalloproteinase GP63 (1995). Biochim Biophys Acta (BBA)/Protein Struct Mol. 1253(2):199–207. DOI:10.1016/0167-4838(95)00155-5

69. Feliciano GT, Da Silva AJR. Unravelling the reaction mechanism of matrix metalloproteinase 3 using QM/MM calculations (2015). J Mol Struct. 1091:125–32. DOI:10.1016/j.molstruc.2015.02.079

70. WE K, KH J, JH M Z W. Degradation of extracellular matrix by larvae of </i>Schistosoma mansoni</i>. II. Degradation by newly transformed and developing schistosomula. (1983). Lab Invest. 49(2):201–7.

71. Bruce JI, Pezzlo F, McCarty JE, Yajima Y. Migration of Schistosoma mansoni through mouse tissue. Ultrastructure of host tissue and integument of migrating larva following cercarial penetration. (1970). Am J Trop Med Hyg. 19(6):959–81. DOI:10.4269/ajtmh.1970.19.959

72. Breitkreutz D, Koxholt I, Thiemann K, Nischt R. Skin basement membrane: The foundation of epidermal integrity - BM functions and diverse roles of bridging molecules nidogen and perlecan (2013). Biomed Res Int. DOI:10.1155/2013/179784

73. Smith LT, Holbrook KA, Byers PH. Structure of the dermal matrix during development and in the adult (1982). J Invest Dermatol. 79(Suppl. 1):93–104. DOI:10.1038/jid.1982.19

74. Marikovsky M, Levi-Schaffer F, Arnon R, Fishelson Z. Schistosoma mansoni: Killing of transformed schistosomula by the alternative pathway of human complement (1986). Exp Parasitol. 61(1):86–94. DOI:10.1016/0014-4894(86)90138-4

75. Dessein A, Samuelson JC, Butterworth AE, Hogan M, Sherry BA, Vadas MA, et al. Immune evasion by Schistosoma mansoni: Loss of susceptibility to antibody or complement-dependent eosinophil attack by schistosomula cultured in medium free of macromolecules (1981). Parasitology. 82(3):357–74. DOI:10.1017/S0031182000066890

76. Auriault C, Ouaissi MA, Torpier G, Eisen H, Capron A. Proteolytic cleavage of IgG bound to the Fc receptor of Schistosoma mansoni schistosomula (1981). Parasite Immunol. 3(1):33–44. DOI:10.1111/j.1365-3024.1981.tb00383.x

77. Pleass RJ, Kusel JR, Woof JM. Cleavage of human IgE mediated by Schistosoma mansoni (2000). Int Arch Allergy Immunol. 121(3):194–204. DOI:10.1159/000024317

78. Bourke CD, Prendergast CT, Sanin DE, Oulton TE, Hall RJ, Mountford AP. Epidermal keratinocytes initiate wound healing and pro-inflammatory immune responses following percutaneous schistosome infection (2015). Int J Parasitol. 45(4):215–24. DOI:10.1016/j.ijpara.2014.11.002

79. Chen L, Rao KVN, He YX, Ramaswamy K. Skin-stage schistosomula of Schistosoma mansoni produce an apoptosis-inducing factor that can cause apoptosis of T cells (2002). J Biol Chem. 277(37):34329–35. DOI:10.1074/jbc.M201344200

80. Sanin DE, Mountford AP. Sm16, a major component of Schistosoma mansoni cercarial excretory/secretory products, prevents macrophage classical activation and delays antigen processing (2015). Parasites and Vectors. 8(1):1. DOI:10.1186/s13071-014-0608-1

81. Ramaswamy K, Salafsky B, Lykken M, Shibuya T. Modulation of IL-1alpha, IL-1beta and IL-1RA production in human keratinocytes by schistosomulae of Schistosoma mansoni (1995). Immunol Infect Dis. 5(2):100–7.

82. Ramaswamy K, Potluri S, Ramaswamy P, … AE-… 9th IC, 1998 U. Immune evasion by Schistosoma mansoni: characterization of Sm 16.8 an anti-inflammatory protein produced by the skin stage schistosomulum (1998). Proc 9th Int Conf Parasitol. :597–603.

83. Holmfeldt P, Brännström K, Sellin ME, Segerman B, Carlsson SR, Gullberg M. The Schistosoma mansoni protein Sm16/SmSLP/SmSPO-1 is a membrane-binding protein that lacks the proposed microtubule-regulatory activity (2007). Mol Biochem Parasitol. 156(2):225–34. DOI:10.1016/j.molbiopara.2007.08.006

84. Ramaswamy K, Kumar P, He Y-X. A Role for Parasite-Induced PGE 2 in IL-10-Mediated Host Immunoregulation by Skin Stage Schistosomula of Schistosoma mansoni (2000). J Immunol. 165(8):4567–74. DOI:10.4049/jimmunol.165.8.4567

85. Hervé M, Angeli V, Pinzar E, Wintjens R, Faveeuw C, Narumiya S, et al. Pivotal roles of the parasite PGD2 synthase and of the host D prostanoid receptor 1 in schistosome immune evasion (2003). Eur J Immunol. 33(10):2764–72. DOI:10.1002/eji.200324143

86. McKean JR, Anwar ARE, Kay AB. Schistosoma mansoni: Complement and antibody damage, mediated by human eosinophils and neutrophils, in killing schistosomula in vitro (1981). Exp Parasitol. 51(3):307–17. DOI:10.1016/0014-4894(81)90118-1

87. Kitaura M, Nakajima T, Imai T, Harada S, Combadiere C, Tiffany HL, et al. Molecular cloning of human eotaxin, an eosinophil-selective CC chemokine, and identification of a specific eosinophil eotaxin receptor, CC chemokine receptor 3 (1996). J Biol Chem. 271(13):7725–30. DOI:10.1074/jbc.271.13.7725

88. Wechsler ME, Munitz A, Ackerman SJ, Drake MG, Jackson DJ, Wardlaw AJ, et al. Eosinophils in Health and Disease: A State-of-the-Art Review (2021). Vol. 96, Mayo Clinic Proceedings. Elsevier; 2021. p. 2694–707. DOI:10.1016/j.mayocp.2021.04.025

89. Matisz CE, Leung G, Reyes JL, Wang A, Sharkey KA, McKay DM. Andoptive transfer of helminth antigen-pulsed dendritic cells protects against the development of experimental colitis in mice (2015). Eur J Immunol. 45(11):3126–39. DOI:10.1002/eji.201545579

90. Maizels RM, Balic A, Gomez-Escobar N, Nair M, Taylor MD, Allen JE. Helminth parasites - Masters of regulation (2004). Vol. 201, Immunological Reviews. 2004. p. 89–116. DOI:10.1111/j.0105-2896.2004.00191.x

91. Rook GAW. Review series on helminths, immune modulation and the hygiene hypothesis: The broader implications of the hygiene hypothesis [Internet] (2009). Vol. 126, Immunology. John Wiley & Sons, Ltd; 2009 [cited 2022 Sep 7]. p. 3–11. DOI:10.1111/j.1365-2567.2008.03007.x

92. Maizels RM, Yazdanbakhsh M. Immune regulation by helminth parasites: Cellular and molecular mechanisms [Internet] (2003). Vol. 3, Nature Reviews Immunology. Nature Publishing Group; 2003 [cited 2022 Sep 7]. p. 733–44. DOI:10.1038/nri1183

93. Horák P, Mikeš L, Lichtenbergová L, Skála V, Soldánová M, Brant SV. Avian schistosomes and outbreaks of cercarial dermatitis (2015). Clin Microbiol Rev. 28(1):165–90. DOI:10.1128/CMR.00043-14

